# Distinct processing of visual motion by transcriptomic cell types in the superior colliculus

**DOI:** 10.64898/2026.10.30.762788

**Authors:** Chen Chen, Yuanming Liu, Delaney G. Eaves, Christina I. Lagas, Jiale Ren, Arda Kipcak, Hui Chen, Alev Erisir, Jianhua Cang

**Affiliations:** Department of Psychology, University of Virginia, Charlottesville, VA 22904, USA; Department of Biology, University of Virginia, Charlottesville, VA 22904, USA; Neuroscience Graduate Program, University of Virginia, Charlottesville, VA 22904, USA

## Abstract

How the brain organizes visual processing across distinct neuron types is a fundamental yet unresolved question. Here we study how visual motion is processed by transcriptomically defined cell types in the superior colliculus (SC). We generated or identified transgenic mouse lines to target three transcriptomic types of neurons in the superficial SC, the *Cbln4+*, *Vip+*, and *Grp+* neurons. We find that each of these cell types encodes distinct ranges of stimulus direction, speed, size, and spatial and temporal features. In response to ecologically relevant visual flows, the three cell types occupy partitional spaces in the encoding manifold. The three cell types also show unique morphology and connectivity, including their retinal and intracollicular input and their projection patterns. Together, these results reveal a genetic logic for visual motion processing in the SC, where transcriptomic cell types are likely the basic computational units with their specific feature selectivity, morphology, and input-output connectivity.

## Introduction

Since the early days of neuroscience, scientists have tried to parse the brain into cell types. Early studies classified neurons based on morphology and later ones based on function. More recently, single-cell genomics has classified neuronal subtypes by their transcriptome^1–3^. These studies have allowed unbiased classification of cell types at a greater level of granularity. However, the link between transcriptomic cell types and functional and morphological properties is not a straightforward matter. In the visual system, for example, a clear correspondence is seen between transcriptomic types and functional properties for retinal ganglion cells^4^, but not for neurons in the zebrafish optic tectum or inhibitory neurons in the mouse visual cortex^5,6^. The link between cell types defined by different modalities therefore remains unclear; yet understanding it is a critical step for studying how the nervous system functions, develops, and evolves.

Here we address this fundamental issue in the superior colliculus (SC). The SC is a major hub for visual motion processing, particularly in visually guided innate behaviors^7–11^. The mouse SC is known to respond to the visual motion caused by running prey or looming predators and, in turn, drive proper motor actions^12–15^. In addition, it has been shown in both primates and rodents that the SC plays a key role in many higher cognitive functions that are usually associated with the cerebral cortex, such as visual spatial attention and perceptual decision-making^16–20^. Despite these extensive studies, however, it remains largely unclear how visual motion is encoded and transformed by the diverse types of neurons in the SC.

Diverse visual responses are seen in the SC, including motion-sensitive neurons tuned to specific features, such as motion direction and speed^21,22^. Early studies revealed the broad anatomical organization of these response properties. For example, the superficial layers of the SC (sSC) are purely visual and receive direct input from the retina^23–26^. Later studies attempted to correlate visual responses of SC neurons with their somatodendritic morphology^22,27–31^. These studies reported differences in response properties between morphological cell types, but no unique corraespondence was observed between morphology and visual functions. Recent studies using single-cell genomics have revealed far more neuronal subtypes in the SC than previously recognized^32–35^. It was shown that the mouse sSC contains ∼30 transcriptomic neuron types^32^. These types are remarkably conserved across species, from mice to tree shrews to humans^36^, suggesting that they are likely the organizing units of SC circuits and computation. These studies thus raise a fundamental question and exciting possibility regarding the relationship between function and cell types: do transcriptomic cell types encode distinct visual motion features and exhibit unique anatomical properties?

To answer this question, we used a genetic approach to access three transcriptomic neuron types in the SC. We found each of the cell types prefers a distinct range of stimulus direction, speed, size, and spatial and temporal frequency. In response to an ecologically relevant stimulus called “flows”, the three cell types occupy partitional spaces of the encoding manifold and show multiplexed encoding and unique visual motion processing. The three cell types also exhibit distinct morphology, retinal and intracollicular inputs, and projection patterns. Together, these results reveal a genetic logic for motion processing in the SC, where individual transcriptomic cell types receive specific synaptic inputs, encode unique combinations of visual features, and process the information locally or send to different downstream targets. These findings will not only profoundly shape our understanding of SC function, circuitry, and development, but also provide valuable information to the fundamental task of defining cell types in the nervous system.

## Results

### Genetically accessing transcriptomic cell types in the SC

We used a genetic approach to gain access to transcriptomically defined cell types to study how they encode and transform visual information (Fig. 1a). We focused on the superficial SC as it is enriched with neurons that are motion sensitive, including direction selective (DS) neurons^37^. First, *Cbln4 (Cerebellin 4)* is a marker gene for a subtype of inhibitory neurons in the sSC (Fig. 1b, cluster n27 in our single-cell RNA-seq atlas)^32^. Given the abundance and diversity of inhibitory neurons in the sSC (∼60% of all neurons and consisting of 18 transcriptome subtypes), we generated a transgenic mouse line using CRISPR/Cas9 approach to target the *Cbln4+* cells (Cbln4-IRES-Cre, Fig. 1c). We performed RNA Fluorescence In Situ Hybridization (FISH) and confirmed colocalization of the *Cbln4* and Cre transcripts (Fig. 1d). We further tested this line by injecting Cre-dependent AAV to express tdTomato (AAV-hSyn-FLEX-tdTomato) in Cre+ cells and quantified the colocalization between tdTomato and *Cbln4* transcripts (Fig. 1e). The vast majority (82.5% ± 1.2%) of *Cre+* cells in the superficial 150 μm of the SC were found to be *Cbln4*+ (“specificity”), and 71.8% ± 5.3% of *Cbln4*+ cells were tdTomato+ (“sensitivity”, Fig. 1f), confirming the validity of this Cre line.

**Fig. 1:**
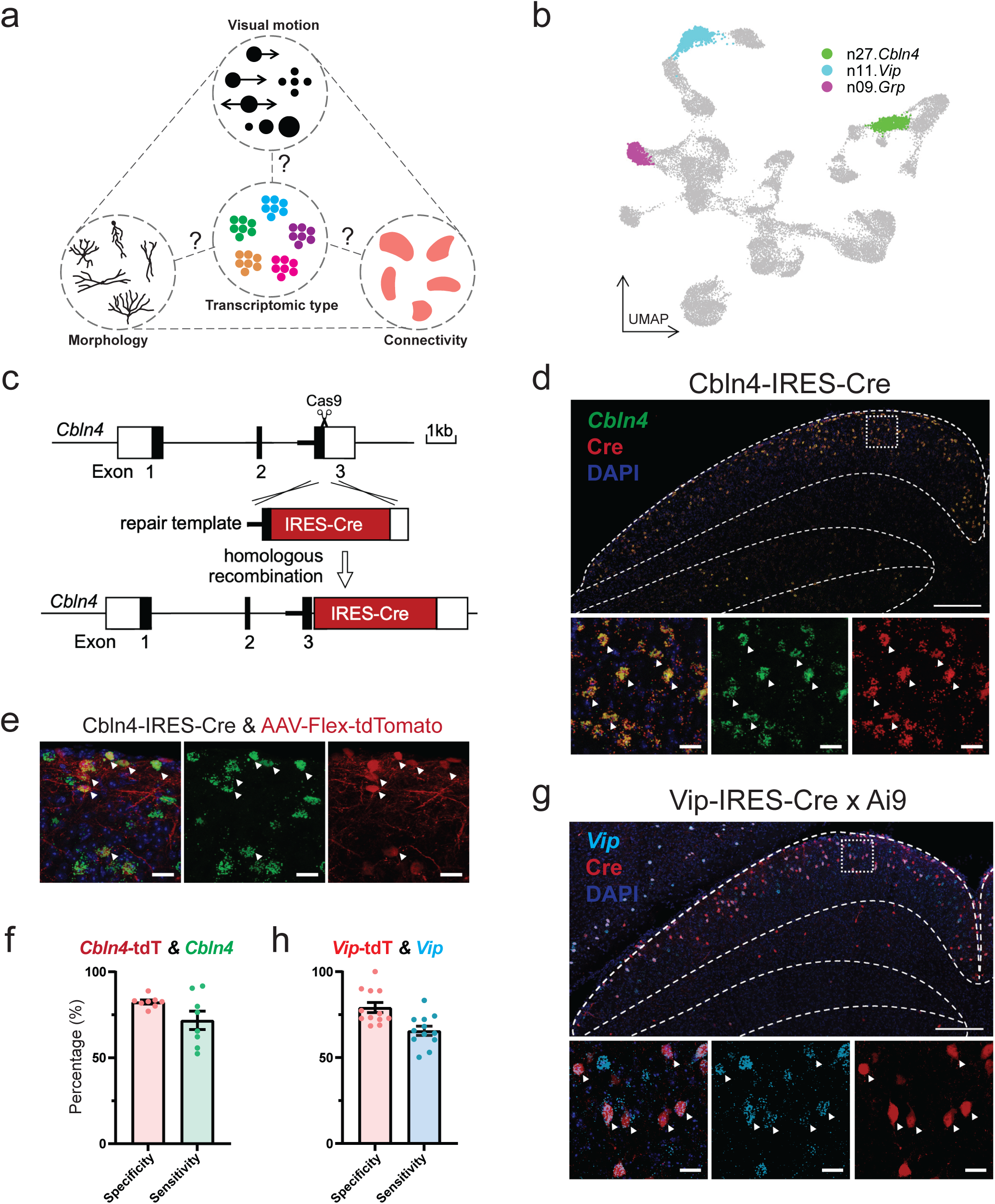
Genetically accessing distinct transcriptomic cell types with transgenic mouse lines a, Schematic of studying the function, morphology and connectivity of transcriptomically defined cell types. **b,** UMAP plot illustrating the expression of selected marker genes in distinct clusters. **c,** Design of the Cbln4-IRES-Cre mouse line using CRISPR-Cas9. **d,** RNA FISH of *Cbln4* and Cre expression in the sSC of Cbln4-IRES-Cre mice. The area of white square in the top panel is shown at a higher magnification with merged and individual channels in the bottom panels. Example colocalized cells are marked by white arrows. Scale bars: 200 μm (top) and 20 μm (bottom). **e,** RNA FISH of *Cbln4* expression and Cre-dependent tdTomato expression in the sSC of Cbln4-IRES-Cre mice. Example colocalized cells are marked by white arrows. Scale bar: 20 μm. **f,** Bar plot showing the specificity and sensitivity of tdTomato labeled Cre expression of Cbln4-IRES-Cre line with *Cbln4* expression. Mean ± SEM. n = 8 images, 4 mice. **g,** RNA FISH of *Vip* expression in the sSC of Vip-IRES-Cre x Ai9 mice. The area of white square in the top panel is shown at a higher magnification with merged and individual channels at the bottom. Example colocalized cells are marked by white arrows. Scale bars: 200 μm (top) and 20 μm (bottom). **h,** The specificity and sensitivity of Vip-IRES-Cre line with *Vip* expression. Mean ± SEM. n = 12 images, 3 mice.

Second, we chose to study a subtype of excitatory neurons that are located in the same lamina as *Cbln4+* cells. This type expresses the marker gene *V*i*p* (*Vasoactive Intestinal Peptide*, Cluster n11; Fig. 1b). We examined a widely used Vip-Cre line^38^ but found that it did not faithfully label *Vip+* cells in the SC^32^. We therefore studied a different Vip-IRES-Cre mouse line^39^. We crossed this line with a reporter line Ai9 (tdTomato) to visualize Cre expression and used RNA FISH to stain the *Vip* transcript (Fig. 1g). Both Cre+ and *Vip*+ cells were mainly located in the upper SGS (*stratum griseum superficiale*) as expected, with a specificity of 79.1% ± 2.9% and sensitivity of 65.7% ± 2.7% (Fig. 1h).

Third, we studied *Grp*+ (Gastrin Releasing Peptide) cells, which are the excitatory Cluster n09 in our single cell sequencing atlas (Fig. 1b) and located slightly below *Cbln4+* and *Vip+* cells in the lower SGS. To do this, we used Grp-KH288-Cre mice, which we had previously demonstrated to be a faithful line for this cell type^40^. These 3 mouse lines thus allowed us to study the visual function, morphology, and input/output connectivity of these molecular cell types in the SC.

### *Cbln4+, Vip+ and Grp+* cells display distinct direction selectivity to drifting gratings

We directly measured the neuronal activity of each cell type using two photon imaging of Cre-dependent expression of calcium indicators (AAV-hSyn-FLEX-GCaMP6s) in awake head-fixed mice (Fig. 2a,b). We first mapped the receptive field of individual neurons in each imaging session using flashing squares. All three types of cells responded strongly (Extended Data Fig.1) and showed similar receptive field structure and retinotopic progression (Extended Data Fig.2).

**Fig. 2:**
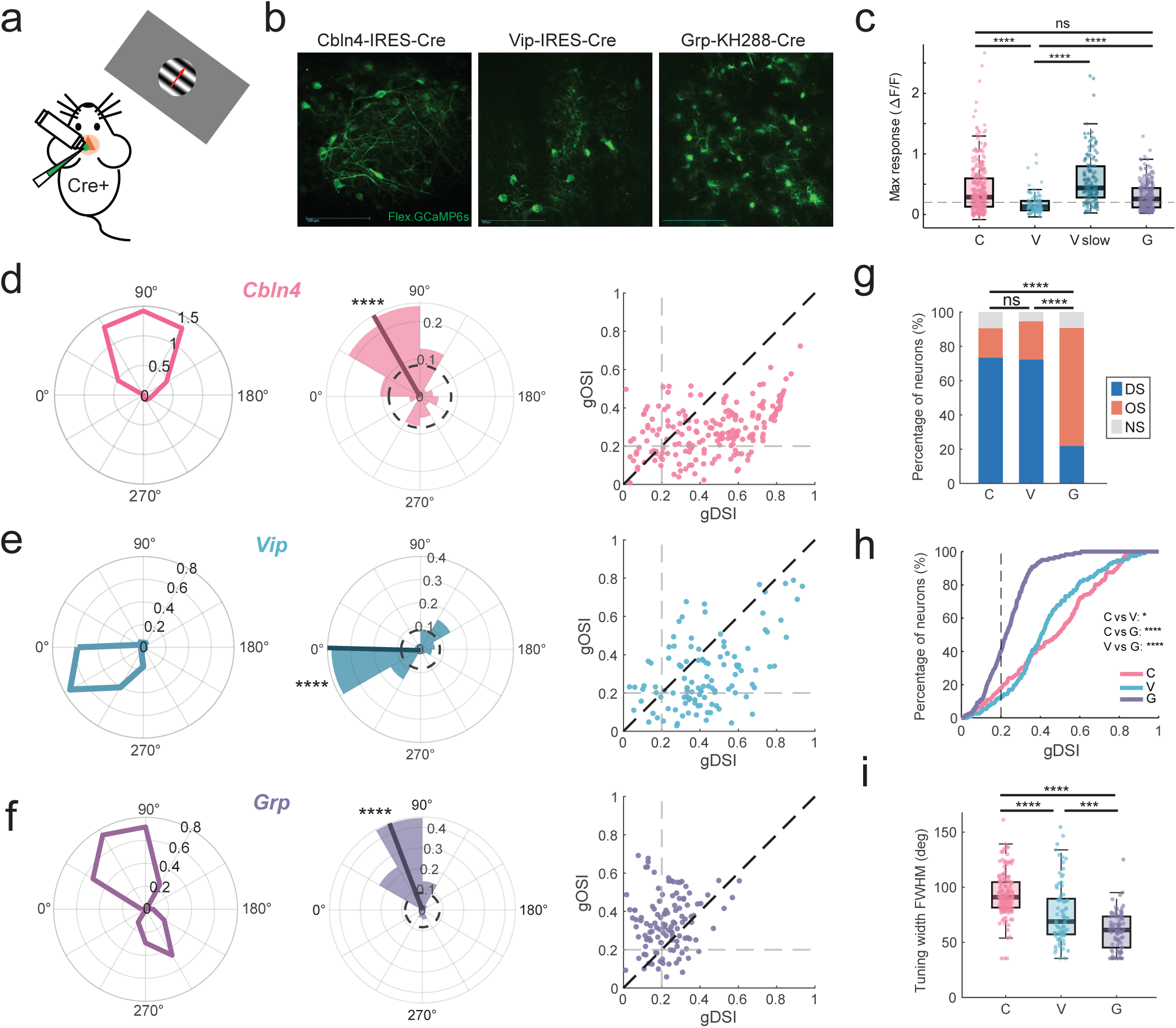
*Cbln4+, Vip+ and Grp+* cells display distinct direction selectivity to drifting gratings. **a**, Schematic of 2-photon recording of the SC in response to drifting gratings. **b,** Example 2-p images showing GCaMP6S expression in the 3 cell types. Scale bars: 100 μm. **c,** Box plot showing the max response of all recorded neurons. C (*Cbln4*), V (*Vip*), and G (*Grp*) label the corresponding cells recorded in response to drifting gratings (SF = 0.16 cpd, TF = 2 Hz, speed of 25 °/sec). V-slow is the *Vip*+ cells recorded to slow gratings (SF = 0.16 cpd, TF = 0.8 Hz, speed of 5 °/sec; and SF = 0.08 cpd, TF = 0.5 Hz, speed of 6.25 °/sec). C: n = 323 cells, 5 mice; V: n = 88 cells, 6 mice; V_slow: n = 145 cells, 11 mice. G: n = 232 cells, 9 mice. Group compared by Kruskal-Wallis test (H(3) = 92.86, p < 0.001), with Dunn’s post-hoc test (Holm-corrected). ****p < 0.0001; ns, not significant. C vs G was not significant (p = 0.079); all other pairwise comparisons were significant (p < 0.0001). **d,** Tuning properties for *Cbln4+* cells. Left: Polar plot showing example tuning curves to drifting gratings. r axis: ΔF/F. Middle: Rose histogram showing the distribution of the preferred direction. Only responsive and selective cells were included (gDSI ≥ 0.2 and gDSI > gOSI; n = 141 cells, 5 mice). Solid line indicates the circular mean; dashed circle indicates an even distribution for reference. Rayleigh test, p < 0.0001. Axial Rayleigh test, p < 0.0001. Right: Scatter plot of gDSI vs gOSI for all responsive cells (n = 192 cells, 5 mice). **e,** Tuning properties for *Vip+* cells, as in **d** Middle: gDSI ≥ 0.2 and gDSI > gOSI; n = 81 cells, 11 mice. Rayleigh test, p < 0.0001. Axial Rayleigh test, p < 0.0001. Right: n = 112 cells, 11 mice. **f,** Tuning properties for *Grp+* cells, as in **d** Middle: gDSI ≥ 0.2 or gOSI ≥ 0.2; n = 108 cells, 9 mice. Rayleigh test, p = 0.0029. Axial Rayleigh test, p < 0.000. Right: n = 119 cells, 9 mice. **g,** Bar plot showing the percentage of direction-selective (DS), orientation selective (OS), and non-selective (NS) cells. C: n = 192 cells, 5 mice; V: n = 112 cells, 11 mice; G: n = 119 cells, 9 mice. Pairwise Chi-squared test with Holm corrected (C vs V: p = 0.296; C vs G, V vs G: p < 0.0001). **h,** Cumulative percentage of gDSI for the 3 cell types of responsive cells. Pairwise Kolmogorov-Smirnov test with Holm corrected (C vs V: p = 0.0176; C vs G, V vs G: p < 0.0001). **i,** Box plot of the tuning width of the 3 cell types. Only selective and well fitted cells were included (gDSI ≥ 0.2 or gOSI ≥ 0.2; with r^2^ > 0.7). Group compared by Kruskal-Wallis test (H(2) = 114.6, p < 0.001), with Dunn’s post-hoc test (Holm-corrected). C vs V, C vs G: p < 0.0001; V vs G: p = 0.00076.

Next, we studied direction selectivity using drifting gratings of the same parameters as in previous studies: spatial frequency (SF) of 0.08 cycles per degree (cpd) and temporal frequency (TF) of 2 Hz (for a speed of 25 °/sec). Most *Cbln4*+ (59.4%) and *Grp*+ (51.3%) cells were responsive to this stimulus (see Methods for details of responsiveness criteria, Extended Data Fig.1). In contrast, only 23.9% of *Vip*+ cells were responsive (Extended Data Fig.1). Even for the responsive ones, *Vip*+ cells showed much weaker responses than *Cbln4*+ and *Grp*+ cells (Fig. 2c, Extended Data Fig. 3a). We then explored the possibility that *Vip*+ cells might prefer a different drifting speed and found them in fact responded much more strongly to slower gratings (Fig. 2c, Extended Data Fig. 3b, c). We used two sets of slower gratings (SF = 0.16 cpd, TF = 0.8 Hz, speed of 5 °/sec; and SF = 0.08 cpd, TF = 0.5 Hz, speed of 6.25 °/sec) to determine direction selectivity of *Vip*+ cells and combined them in the following analysis.

*Cbln4+, Vip+ and Grp+* cells all showed highly selective responses to gratings, but with distinct profiles (Fig. 2d-f). We calculated direction and orientation selectivity index (gDSI and gOSI, see methods for details) for each neuron. For *Cbln4+* cells, the vast majority of responsive ones were highly DS (73.4% of cells with gDSI ≥ 0.2 & gDSI > gOSI) and showed stereotypical DS tuning curves (Fig. 2d). Strikingly, most of these cells preferred similar directions, instead of evenly distributed over the entire possible range (Fig. 2d). Nearly half of the cells (46.1%) preferred directions between 30° and 90° (Fig. 2d; 0° for nasal and 90° for upward direction on the stimulus monitor), showing a population preference near upward motion (circular mean 60.2°, R = 0.440). Similarly, *Vip*+ cells were also highly DS in response to the slow gratings (72.3% with gDSI ≥ 0.2 & gDSI > gOSI). However, unlike *Cbln4+* cells, *Vip+* cells mostly preferred nasally directed motions (circular mean 1.5°, R = 0.267, Fig. 2e). More than half (53.1%) of the cells preferred directions between 300° and 360° (Fig. 2e).

In contrast, *Grp*+ cells showed a different response profile, often with two peaks ∼180° apart in their tuning curves (Fig. 2f). In other words, *Grp*+ cells were mostly selective for stimulus axis or orientation. Interestingly, almost all *Grp*+ cells showed a preference near the vertical axis, often with stronger responses for the upward motion (circular mean 69.2°, R = 0.690, Fig. 2f). Consistent with the two-peaked tuning curves, most *Grp*+ cells had higher gOSI than gDSI (Fig. 2f), and they were classified as orientation/axis selective (Fig. 2g). This is different from *Cbln4+* and *Vip+* cells, most of which had higher gDSI than gOSI and were DS (Fig. 2d-h). Additionally, *Cbln4*+ cells had a broader tuning width (median of 90.9°) than *Vip*+ (median of 68.9°) and *Grp*+ (median = 61.2°) cells (Fig. 2i).

Together, these results demonstrate that the 3 molecular cell types encode distinct visual motion information, each uniquely tuned for a specific motion direction (or axis) and speed.

### Cell-type specific spatial and temporal selectivity, size preference, and speed tuning

We next tested whether the functional difference extends beyond direction selectivity by presenting a battery of visual stimuli to probe other features. First, we systematically varied the spatial and temporal frequency of drifting gratings (0.02 to 0.64 cpd; 0.25 to 8 Hz, Fig. 3a). The gratings were displayed near the preferred direction for each cell type (90° for *Cbln4*+ and *Grp*+; and 330° for *Vip*+ cells) and evoked reliable responses in nearly all the cells (*Cbln4*+ 92.5%, *Vip*+ 100%, *Grp*+ 80.4%; Extended Data Fig.1). *Cbln4*+ cells responded broadly in the SF and TF space, with the strongest responses at the middle SFs (0.08-0.16 cpd, Fig. 3b-e, Extended Data Fig.3d-f). *Vip*+ cells preferred lower SFs than *Cbln4*+ cells (Fig. 3b-d). *Cbln4*+ and *Vip*+ cells both showed preference for low TFs of 0.5-1 Hz (Fig. 3e). In contrast, *Grp*+ cells responded more strongly to higher TFs (Fig. 3b-e), with most cells (77.8%) preferring 4-8Hz.

**Fig. 3:**
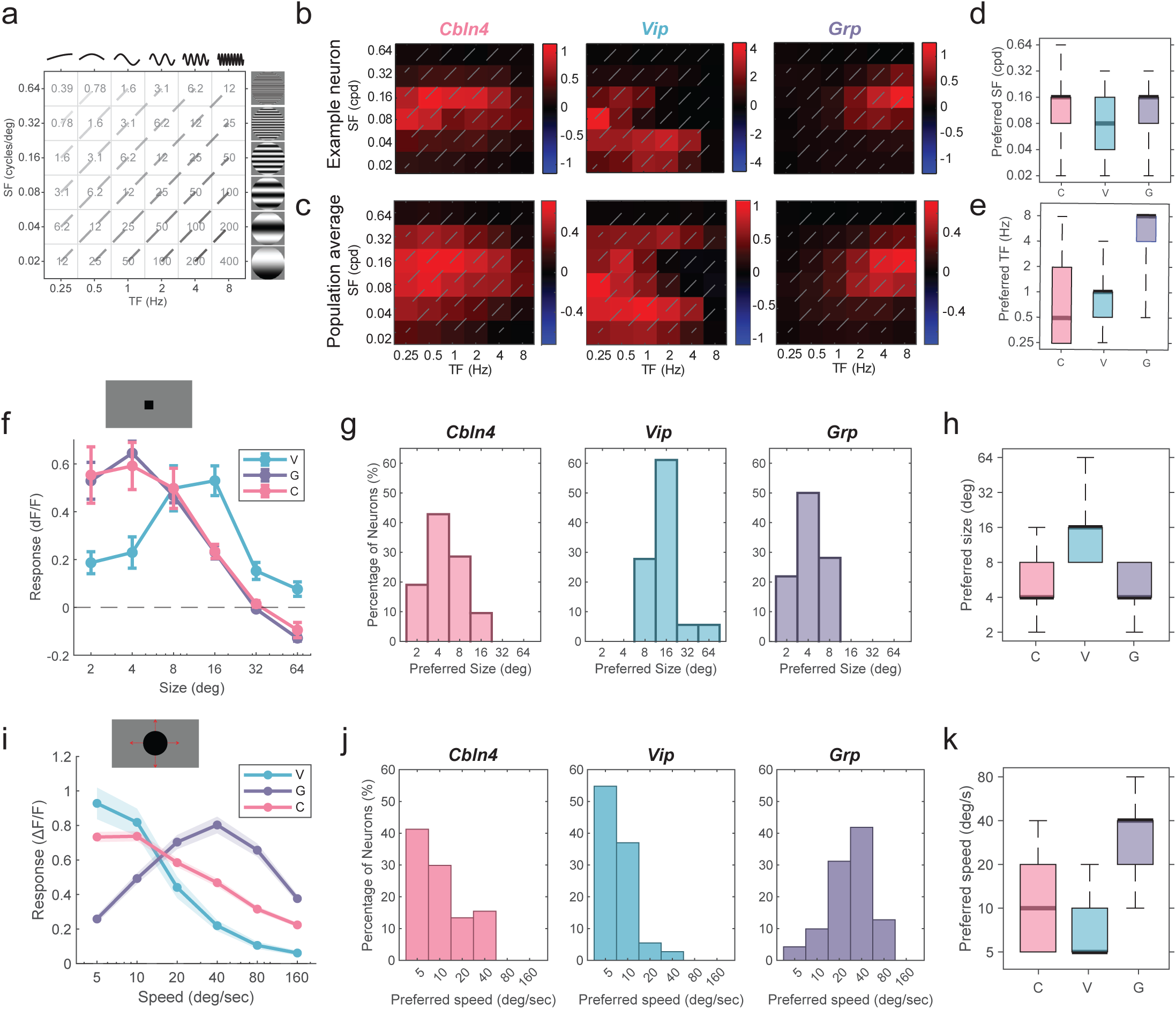
*Cbln4+, Vip+ and Grp+* cells prefer different spatial and temporal frequency, size, and speed. **a,** Schematic of drifting gratings displayed in a series of SF and TF combinations. Dashed lines of different colors indicate the same speed. **b,** Heatmap showing the response magnitude of 3 example cells for different SF-TF combinations (left to right: *Cbln4+, Vip+ and Grp+.* The same order is followed in all panels). **c,** Heatmap showing the average response magnitude of all responsive cells from the 3 cell types. C: n = 172 cells, 4 mice; V: n = 23 cells, 3 mice; G: n = 45 cells, 3 mice. **d-e,** Box plot of the distribution of preferred SF (**d)** and TF (**e)** of the 3 cell types. **f,** Schematic of flashing square (top). Average tuning curve of all responsive cells to flashing squares of different sizes (bottom). C: n = 21 cells, 4 mice. V: n = 18 cells, 8 mice. G: n = 32 cells, 5 mice. Mean ± SEM. **g-h,** Histogram **g** and box plot **h** showing the distribution of preferred size of the 3 cell types. **i,** Schematic of looming (top). Average tuning curve of all responsive cells to looming expanding at different speeds (bottom). C: n = 194 cells, 7 mice. V: n = 73 cells, 6 mice. G: n = 141 cells, 8 mice. Mean ± SEM. **j-k,** Histogram **j** and box plot **k** showing the distribution of preferred speed of the 3 cell types.

Second, we asked whether the 3 molecular cell types encode distinct size information, using a black square of different sizes centered on the receptive field (Fig. 3f, Extended Data Fig.1). Both *Cbln4*+ and *Grp*+ responded more strongly to the smaller sizes (2-8°, strongest to 4° squares), and they were suppressed by the largest size tested (Fig. 3f, g). In contrast, no *Vip*+ cells showed a preference for the small 2° and 4° squares (Fig. 3g). Instead, *Vip*+ cells responded the strongest to 16° squares (Fig. 3f-h).

Next, we studied speed tuning using “looming”, which have been widely used to mimic approaching predators (Fig. 3i and Extended Data Fig.1). *Grp*+ cells responded strongly to looming and showed the highest responses to 40°/sec speed (Fig. 3i-k). *Cbln4*+ and *Vip*+ cells responded much weaker at this speed (Fig. 3i). Instead, they preferred slower ones, especially at 5-10 °/sec (Fig. 3i-k). Particularly, *Vip*+ cells were barely active to the 40 °/sec stimulus (Fig. 3i). Notably, the different speed preferences of the 3 cell types are highly consistent between the looming and grating stimuli (Extended Data Fig. 3g, h).

### *Cbln4+, Vip+ and Grp+* cells occupy partitional spaces in an encoding manifold

Natural visual scenes rarely vary along a single feature at a time. We therefore used an ecologically relevant stimulus called “flows”, in which multiple features vary simultaneously. The flow stimulus consisted of a full field of patterned black dots of a certain size (2-16°, 4 steps) that moved at a certain speed (0-40°/sec, 5 steps), direction (12 directions), and motion jitter (see Methods for details, Fig. 4a). The large number of stimulus conditions allowed us to examine how the 3 cell types are organized in the high-dimensional stimulus space. We embedded flow-evoked responses of all recorded neurons into an encoding manifold^41^. In this approach, neurons with similar response profiles are positioned close to one another, and cell-type identity is not used to construct the manifold. Nevertheless, the 3 types were clearly separated in the first three diffusion coordinates (Fig. 4b). Notably, this organization differed from that of both retinal ganglion cells (RGCs) and primary visual cortex (V1) neurons. RGCs form discrete clusters, whereas V1 neurons are distributed uniformly along a continuum^41^. In contrast, SC cell types occupied distinct regions within a continuous manifold.

**Fig. 4:**
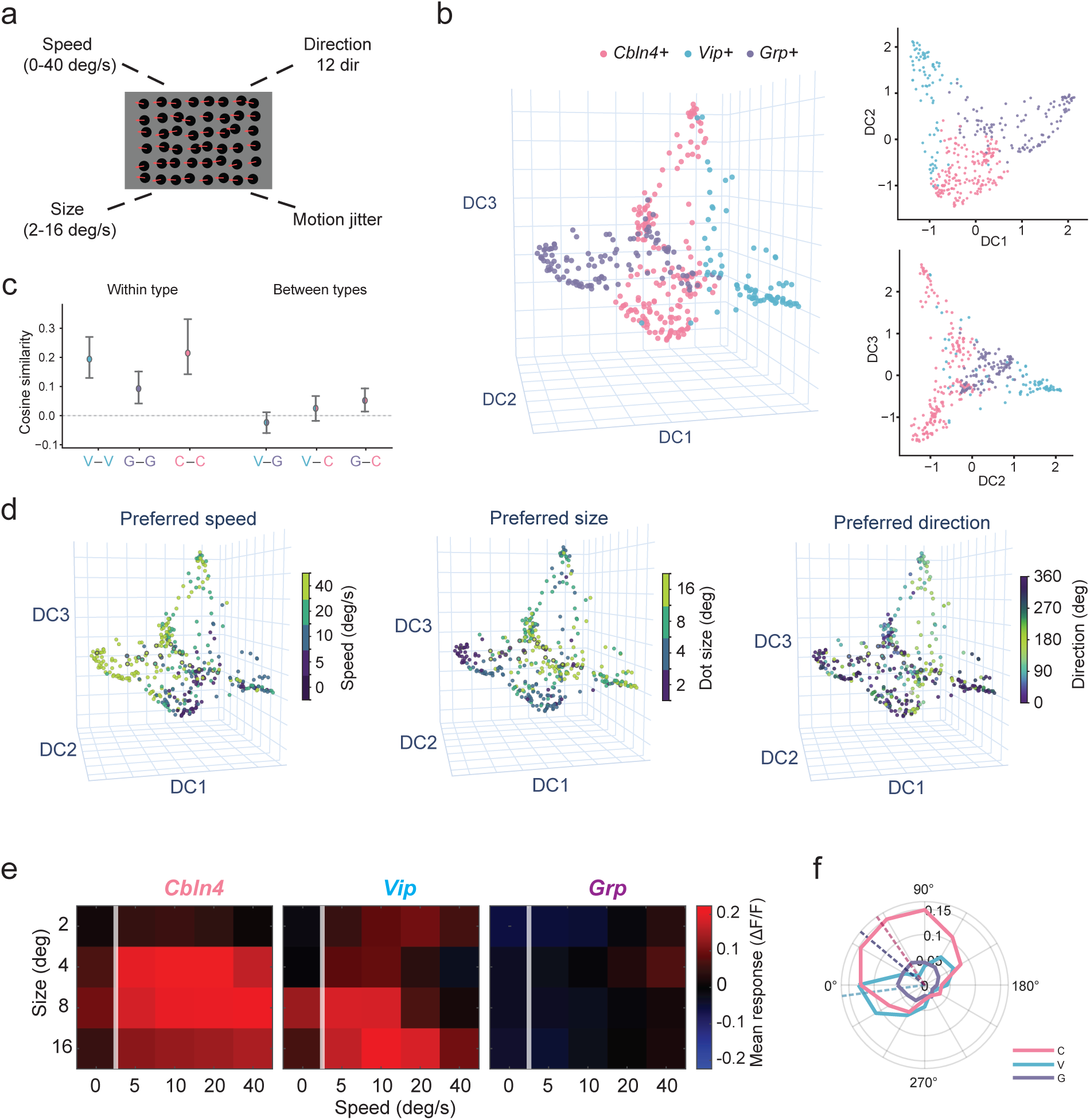
Distinct responses of the 3 cell types to ecologically relevant flow stimulus. **a,** Schematic of the flow stimulus and the tested parameters of direction, speed, and size. **b**, Scatter plot showing encoding manifold of all recorded neurons, projected onto the first three diffusion coordinates (DCs). Left, 3D view; right, 2D projections onto DC1 vs. DC2 (top) and DC2 vs. DC3 (bottom). Each dot represents one neuron, colored by cell type. C: n = 184 cells, 4 mice; V: n = 81 cells, 5 mice; G: n = 115 cells, 5 mice. **c**, Mean cosine similarity of responses to flows between neurons within each cell type (left) and between each pair of cell types (right), averaged over animals. Mean ± SEM. **d**, Encoding manifold as in **b**, with each neuron colored by its preferred speed (left), size (middle), or direction (right) to flows. **e**, Heatmap of the average response of different combinations of stimulus speeds and sizes. All responsive cells from each cell type were included. **f,** Average direction tuning curves averaged over all speeds and sizes. Dashed lines: vector sum of the corresponding tuning curve, C (pink): 55.1°, V (blue): 352.5°, G (purple): 39.8°.

To quantify this separation, we computed cosine similarity between pairs of neurons in the full 196-dimensional space. Similarity was greater between neurons of the same type than between neurons of different types (Fig. 4c). Pairwise Euclidean distances showed the same pattern (Extended Data Fig. 4a, b). This separation was consistent across animals: for individual neurons, their nearest neighbors from other animals were much more frequently of the same transcriptomic type than of a different type (Extended Data Fig. 4c), indicating that the functional differences were driven by cell types instead of individual animals.

We mapped each neuron’s preferred features onto the encoding manifold. Preferred speed, size, and direction all varied progressively on the manifold (Fig. 4d), suggesting that the separation between cell types were driven by the stimulus features and their interactions. To confirm this, we examined the response profiles for each cell type. A speed-size response profile was calculated by averaging responses over all directions (Fig. 4e). *Cbln4*+ cells responded broadly over the tested speeds (compared to the stationary dots, 0°/sec; Fig. 4e) at the middle sizes (4-8°); whereas *Vip*+ cells showed stronger responses to slower speeds (5-10°/sec) and larger sizes (8-16°). In contrast, *Grp*+ cells were suppressed by the full-field static dots (0°/sec) and slower speeds (5-10°/sec; Fig. 4e). Their strongest response was to 4° dots moving at 40°/sec, consistent with the size and speed preference in the previous section (Fig. 3h, k). However, the weak responses of *Grp*+ cells (Extended Data Fig. 4d) indicate that they are unlikely to encode full-field flows.

Both *Cbln4*+ and *Vip*+ cells showed high gDSI for flow directions (*Cbln4*+ 0.440 ± 0.019; *Vip*+ 0.454 ± 0.027, *Grp*+ 0.296 ± 0.025, Extended Data Fig. 4e-g). Strikingly, the average of the raw tuning curves (i.e., without any shift) for each type showed a similar direction preference as that in response to drifting gratings (*Cbln4*+ cells 55.1°, and *Vip*+ cells 352.5°, Fig. 4f and Extended Data Fig. 4h, i). These results thus confirm robust and reliable direction selectivity for *Cbln4*+ and *Vip*+ cells, under both flows and drifting gratings.

Together, these results reveal that the three cell types respond to visual flows with different tuning profiles, with each type encoding a particular combination of stimulus direction, size, and speed. Consequently, they occupy distinct spaces in the encoding manifold.

### *Cbln4+, Vip+ and Grp+* cells display distinct morphology and receive unique sets of inputs

The SC consists of neurons of different morphology. The Grp-KH288-Cre line was in fact used to target the narrow field vertical cells^29^. We thus studied whether the transcriptomically defined *Cbln4*+ and *Vip*+ cell types, as well as *Grp*+ cells, display unique anatomical features that might contribute to the newly discovered functional differences.

First, we examined the morphology of individual neurons by injecting Cre-dependent AAV expressing tdTomato (AAV-CAG-FLEX-TdTomato; Fig. 5a). We reconstructed 20-23 cells in each line, capturing the soma shape, dendritic structure and spread of each cell (See details in Methods), taking care to include specific sublayers where the imaging studies were performed (upper SGS for *Cbln4*+ and *Vip*+ cells and lower SGS for *Grp*+ cells, Fig. 5a, Extended Data Fig. 5a, b). The *Cbln4*+ cells displayed fusiform soma shape and bipolar dendrites that were oriented parallel to the SC surface. Dendrites characteristically bifurcated once close to the soma and extended to span horizontally (mean=285.6 µm, n=20) (Fig. 5b-d). *Vip*+ cells had round soma and multipolar dendrites; they displayed the smallest arbor size (Fig. 5b, c and Extended Data Fig. 5c). *Grp*+ cells were also fusiform and bipolar, but these were oriented perpendicular to the surface (Fig. 5b, d), displaying smaller horizontal spread particularly closer to the soma (Fig. 5c). The qualitative morphological features of *Cbln4+*, *Vip+* and *Grp+* cells therefore showed distinct morphology, consistent with horizontal, stellate and narrow field cells, respectively.

**Fig. 5:**
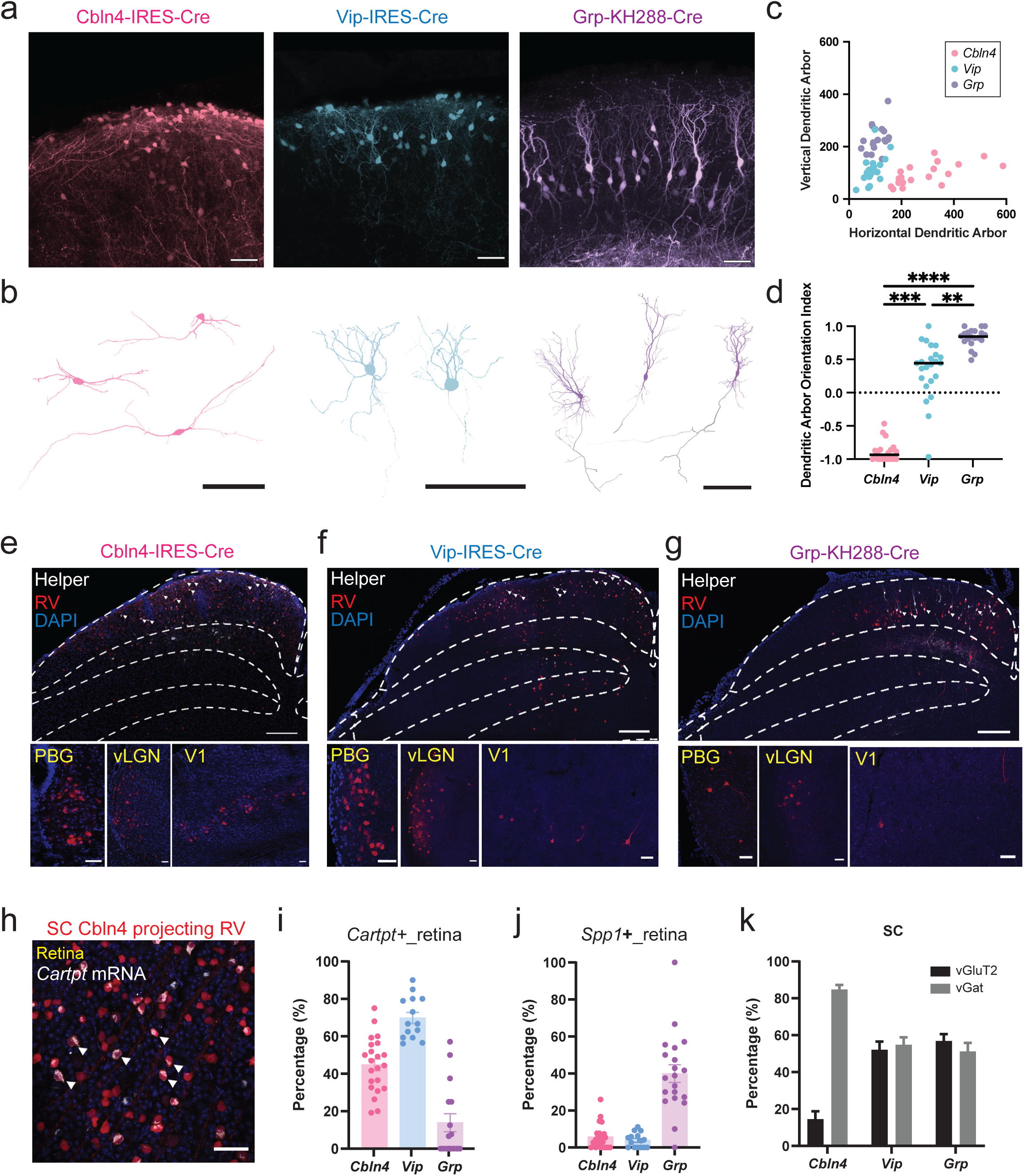
Cell-type specific morphology and input. **a,** Example confocal images showing Cre-dependent tdTomato expression for the 3 cell types in the sSC. Pseudo-colored by the corresponding group colors. Scale bars: 50 μm. **b,** 3D reconstructed morphology of example cells. Gray color labels axons. Scale bars: 100 μm. **c,** Scatter plot presenting the length of horizontal (x axis) and vertical (y axis) dendritic arbor of individual cells. C: n = 20 cells, 3 mice; V: n = 23 cells, 3 mice; G: n = 20 cells, 3 mice. **d,** Quantification of dendritic arbor orientation index. 1 for vertically oriented and -1 for horizontally oriented. Bars label the median. Group compared by Kruskal-Wallis test (H(2) = 49.19, p < 0.001), with Dunn’s post-hoc test (Holm-corrected). C vs V: p = 0.0002, C vs G: p < 0.0001, V vs G: p = 0.0031. **e-g,** Confocal images of rabies tracing in the 3 mouse lines. Upper row: viral injection in the SC, helper virus in white, rabies virus in red, and example starter cells marked in white arrows. Lower row: monosynaptic rabies expressing cells in example brain areas. Scale bars: 200 μm (top) 50 μm (bottom). **h,** Example confocal image showing immunohistochemistry staining of *Cartpt* in the retina and rabies expressing retinal ganglion cells (RGCs) for *Cbln4+* starter cells. White arrow: co-localized example cells. Scale bar: 50 μm. **i,** Quantification of rabies+ RGCs with *Cartpt* expression. C: n = 22 images, 2 mice; V: n = 14 images, 2 mice; G: 16 images, 2 mice. Mean ± SEM. **j,** Quantification of rabies+ RGCs with *Spp1* expression. C: n = 22 images, 2 mice; V: n = 15 images, 3 mice; G: 20 images, 3 mice. Mean ± SEM. **k,** Quantification of sSC non-starter rabies+ cells with vGAT (C: n = 12 images, 2 mice; V: n = 16 images, 4 mice; G: 16 images, 4 mice) or vGluT2 (C: n = 9 images, 2 mice; V: n = 16 images, 4 mice; G: 16 images, 4 mice) expression. Mean ± SEM.

Several differences among the three neuron populations were noted: while both *Vip*+ and *Grp*+ cells had axons that ramify in sSC before coursing ventrally, *Cbln4*+ cells were unique in that an axon emergence point was not observed in any of the 20 reconstructed neurons (Fig. 5b). Similarly, while both *Vip*+ and *Grp*+ cells had spiny dendrites, *Cbln4*+ cell dendrites were smooth (Extended Data Fig. 5d). The normality test based on 7 morphological criteria revealed that *Vip*+ and *Grp*+ cells were morphologically homogeneous, whereas *Cbln4*+ cells were heterogeneous, showing greater variability in many features (Extended Data Fig. 5e).

We next studied what inputs these 3 cell types receive by trans-synaptic rabies tracing. We injected Cre-dependent helper virus (AAV1-hSyn-FLEX-TVA-P2A-GFP-2A-oG) to target each type and then injected monosynaptic retrograde rabies virus (EnvA G-Deleted Rabies-mCherry). Presynaptic neurons expressing mCherry were seen in a number of brain areas related to vision, including the retina, local SC, ventral lateral geniculate nucleus (vLGN), parabigeminal nucleus (PBG), pretectum, and primary visual cortex (V1) for all 3 cell types (Fig. 5e-g, Extended Data Fig.5f). All 3 types receive heavy monosynaptic input from the retina (Fig. 5h, Extended Data Fig. 5g, h). We stained two marker genes to determine the molecular identity of the input cells: *Cartpt* (the cocaine- and amphetamine-regulated transcript neuropeptide) for On-Off direction-selective ganglion cells (DSGCs) and *Spp1* (secreted phosphoprotein 1) for alpha RGCs^43–45^. *Cbln4*+ and *Vip*+ cells received more input from *Cartpt+* than *Spp1+* RGCs (for *Cbln4*+ cells, 44.8 ± 3.1% *Cartpt+* vs. 5.7 ± 1.4% *Spp1+*; for *Vip*+ cells, 69.8 ± 3.0% vs. 3.8 ± 1.0%; Fig. 5i, j), consistent with the observed direction selectivity of *Cbln4*+ and *Vip*+ cells. On the contrary, *Grp*+ cells received more input from *Spp1+* (39.9 ± 4.8%) than *Cartpt+* RGCs (13.8 ± 4.8%; Fig. 5i-j).

Besides the retina, plenty of non-starter rabies+ cells were observed in the sSC locally (Fig. 5h-g and Extended Data Fig. 5i). We stained vesicular GABA transporter (vGAT) and vesicular glutamate transporter 2 (vGluT2) to determine the inhibitory/excitatory identity of these cells. The local input for *Cbln4+* cells was predominantly inhibitory (84.8 ± 2.4% vs. 14.5% ± 4.3% excitatory; Fig. 5k), demonstrating a prominent disinhibitory circuit in the sSC. In contrast, *Vip+* and *Grp+* cells received similar amount of inhibitory and excitatory local SC input (*Vip* 54.9 ± 4.0% vs 52.1 ± 4.4%; *Grp* 51.2 ± 4.6% vs. 56.9 ± 3.7%, Fig. 5k). Together, these results demonstrate that the 3 molecular cell types receive distinct patterns of synaptic inputs, in both retinal direction selectivity and local inhibitory modulation, which likely contribute to their functional differences.

### *Cbln4+, Vip+ and Grp+* cells project to different targets

Finally, we determined the projection pattern of these 3 molecular cell types. Specifically, we focused on the lateral geniculate nucleus (LGN) and the parabigeminal nucleus (PBG), which are the main output of sSC neurons. We injected Cre-dependent AAV to label the Cre+ cells with fluorescence reporters for each line and examined the terminal expression in the LGN and PBG (Fig. 6a). Fluorescent signals were observed in the LGN but not PBG for the Vip-IRES-Cre line (Fig. 6b). Conversely, signals were detected in the PBG but not LGN for the other two lines (Fig. 6b).

**Fig. 6:**
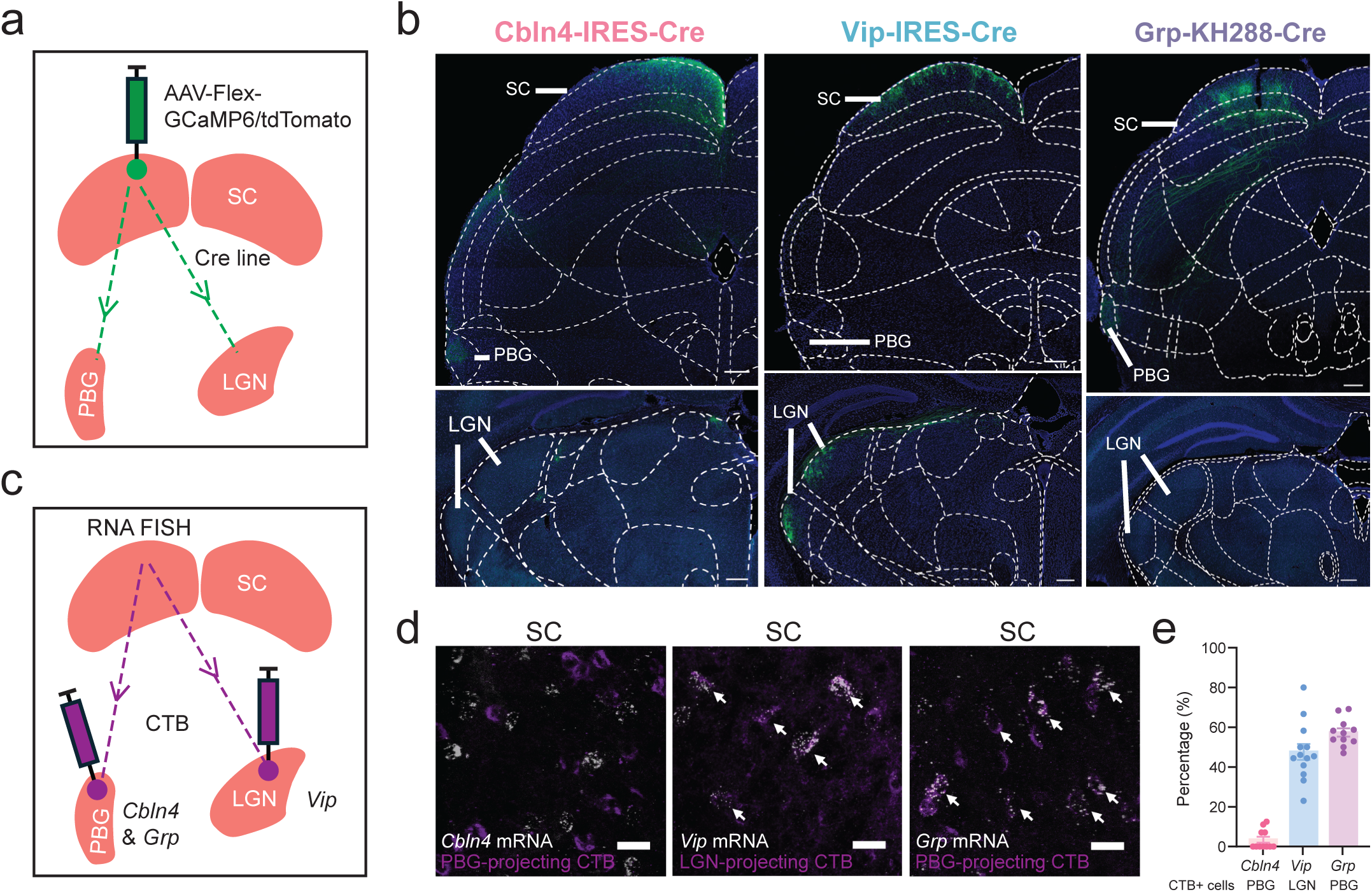
*Cbln4+, Vip+ and Grp+* cells show distinct projection patterns. **a,** Schematic illustrating the method of anterograde tracing. **b,** Confocal images showing Cre-dependent fluorescence expression in the SC (upper row) and terminals in PBG (upper row) and LGN (lower row). Scale bars: 200 μm. **c,** Schematic illustrating the method of retrograde tracing using CTB. **d,** Confocal images showing co-localization of LGN/PBG projecting cells and corresponding marker genes in the sSC. Example colocalized cells marked by white arrows. Scale bars: 20 μm. **e,** Quantification of co-localization of CTB+ cells with the corresponding marker gene in the sSC. C: n = 11 images, 3 mice; V: n = 13 images, 4 mice; G: n =11 images, 3 mice.

To confirm this cell type specific projection pattern, we performed additional experiments using retrograde labeling. We injected Cholera Toxin subunit B (CTB) into either the LGN or PBG and stained the marker genes for corresponding types in the SC (Fig. 6c). For LGN injections, we confirmed the colocalization between CTB+ and *Vip*+ (Fig. 6d). In areas selected for analysis, 47.6 ± 4.0% of the CTB+ cells were *Vip*+ (Fig. 6e). Similarly, for PBG injections, we confirmed that the retrogradely labeled SC neurons were *Grp*+ cells (57.4 ± 2.1% of CTB+ cells were *Grp*+; Fig. 6d, e), consistent with previous reports^29^. However, the colocalization between PBG-projecting CTB+ cells with *Cbln4*+ cells was rare. Only 3.4 ± 1.5% of CTB+ cells were *Cbln4+* (Fig. 6d, e). This indicates that *Cbln4*+ cells, at least those highly expressed in the upper SGS, do not project to the PBG, consistent with our observation that these neurons lack axons. The positive signals observed by anterograde tracing likely originated from low *Cbln4* expression cells in deeper layers or non-specific labelling.

Altogether, our studies reveal that the 3 transcriptomic cell types project to different targets or stay local in the SC, and consequently they play distinct functional roles. *Cbln4*+ cells in the very superficial SC are local inhibitory neurons that prefer small size and upward motion with broad speed preference. These cells show horizontal cell morphology and likely modulate, or even transform, local activity in the SC. *Vip*+ cells are stellate cells that project to the LGN and encode larger objects and slow nasal motion. *Grp*+ cells are narrow field vertical cells that project to the PBG and consistently show preference for small size and fast visual motion. These results thus demonstrate a genetic logic for visual processing in the SC, where transcriptomically defined cell types encode a unique combination of visual features with their distinct morphology, synaptic inputs, and projection patterns (Fig. 7).

**Fig. 7:**
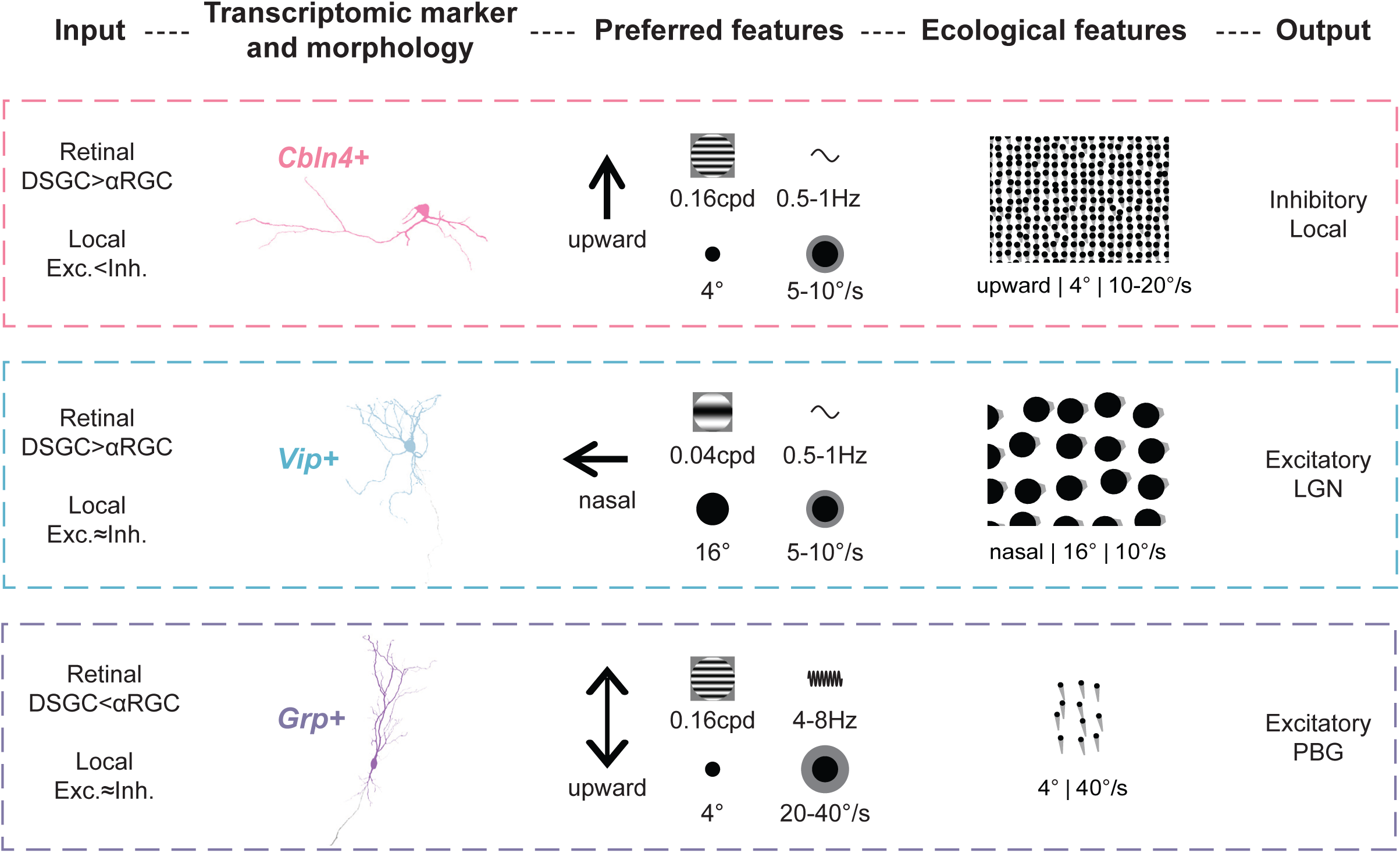
A genetic logic of visual motion processing in the SC. Summary of inputs, morphology, encoded visual features, and outputs for the 3 transcriptomic cell types.

## Discussion

The SC is one of the most studied areas in the brain, extensively investigated across anatomy, function, behavior and evolution. Our study, by targeting transcriptomically defined cell types, represents a major advance in the long-standing pursuit towards a comprehensive understanding of the SC. Conceptually, our results demonstrate that transcriptomic cell types in the SC, with their distinct function, morphology, and connectivity, are the basic units for processing visual motion and mediating visually guided behaviors. Technically, the mouse lines we generated (the Cbln4-IRES-Cre line) or identified (particularly the specific Vip-IRES-Cre line) will facilitate future functional and developmental studies of the SC.

Early studies of the SC revealed its broad anatomical organization, including the layered architecture along the dorsoventral axis and topographic maps within the layers^9,23,46^. These studies also discovered extensive connections that the SC forms with numerous areas throughout the neuraxis^7,23,47^. Later studies classified SC neurons into subtypes based on morphology, physiology, and more recently, transcriptomics. Within the visual layers, several morphological types of neurons are consistently found across species, including horizontal cells, marginal cells, stellate cells, piriform cells, narrow field vertical cells, and wide field vertical cells^7,16,27,46^. Some of the morphological cell types display unique projection patterns; for example, wide field vertical cells project to the pulvinar in all species examined, likely reflecting their conserved roles^16,48–50^. On the other hand, sSC neurons display diverse response properties to visual motion stimuli, including direction selectivity, which is seen in species from fish to rodents to primates^51^. How visual information is processed by the different SC cell types and their circuits thus becomes a critical question.

Studies to correlate morphology with visual response properties were greatly facilitated by the genetic tools available in mice. In a series of studies, Gale and Murphy identified 4 Cre lines to label morphological cell types and investigated their visual functions and projections^28,29,52^. These mouse lines have been very useful for studying how SC morphological cell types contributed to visual processing and behaviors^12,31,53,54^. However, it is important to emphasize that the usefulness of these Cre lines does not necessarily mean that neurons they target are genetically defined cell types. Indeed, the line used to target wide field vertical cells, Ntsr1-GN209-Cre mice, does not reflect endogenous *Ntsr1* expression and also labels non-wide field cells in the deeper SC^40^. More importantly, each morphological cell type may consist of multiple subtypes^55^. This is especially an important concern for targeting horizontal cells using Gad2-Cre, which labels all inhibitory neurons in the SC. We now know that inhibitory neurons are in fact the majority in the sSC and consist of almost 20 transcriptomic subtypes^32^.

In the current study, we aimed to determine whether the cell types defined by single cell transcriptomics display unique function, morphology, and connectivity. We intentionally selected three cell types because of their location near the SC surface, where direction selectivity is high^37^. DS cells in the SC include both excitatory and inhibitory neurons^37^. They are similar in many basic tuning properties, but their synaptic connectivity and ecological roles are largely unknown. By performing *ex vivo* FISH following *in vivo* imaging, our previous study identified *Cbln4+* as a marker gene for inhibitory DS cells^32^. Here, we not only confirmed that *Cbln4+* cells are DS, but discovered that they are tuned to a narrow range of directions (largely upward motion; Fig. 7). *Cbln4+* cells also prefer smaller size and larger SF and show a broad speed tuning. Through their horizontal morphology, *Cbln4+* cells are therefore positioned to mediate lateral inhibition in the SC with their precise direction and spatial information. Interestingly, our retrograde tracing data showed that *Cbln4*+ cells receive prominently inhibitory input within the sSC. In other words, *Cbln4+* cells form part of a disinhibitory circuit, which may mediate direction specific computation, such as encoding motion contrast between stimulus center and surround^56^.

The two excitatory transcriptomic cell types, *Vip*+ and *Grp*+ cells, are tuned to drastically different motion features (Fig. 7). *Vip*+ cells prefer slower motion (5-10 °/s), nasal direction, and larger size (16°); whereas *Grp*+ cells prefer faster motion (20-40 °/s), vertical axis, and smaller size (4°). Notably, *Grp*+ cells are not stereotypical DS as their direction tuning curves often show two peaks along the same axis/orientation; yet they are also different from orientation selective cells found in the cortex as the peak for upward motion is almost always stronger. In addition, *Grp*+ cells are located below *Vip*+ and *Cbln4*+ cells, consistent with the finding that direction selectivity declines with depth^37^. It is also worth emphasizing that the unique tuning properties of the three cell types are remarkably consistent across the many visual stimuli we tested, including the more ecologically relevant visual flows^57^.

More importantly, the striking differences in their feature selectivity indicate that *Vip*+ and *Grp*+ cells play different roles in visual processing and behaviors. Consistent with this idea, it was reported that the speed of a sweeping small black disc (mimicking a cruising predator) affected whether the mouse would freeze or flight: slow sweeps (5 °/sec) led to robust freezing while fast sweeps (> 40 °/sec) increased the probability of flight^58^. On the other hand, the features that *Grp*+ cells are tuned to are similar to those causing approaching behaviors during prey capture (optimal size: ∼5 deg, optimal speed: <50 deg/s)^53,59,60^. Indeed, these cells (i.e., narrow field vertical cells labeled by Grp-KH288-Cre mice) were previously shown to be involved in prey capture^12,53^.

*Vip*+ and *Grp*+ cells also receive different retinal inputs and project to distinct targets, further suggesting their different roles in visual processing. In particular, the SC-to-LGN projection was recently shown to enhance motion selectivity in the LGN^61^ and could subsequently influence V1 responses^62^. With an ascending projection to the LGN, *Vip*+ cells can therefore potentially impact higher cortical functions in visual processing. In contrast, with a descending projection to the PBG, *Grp*+ cells are more likely to be involved in innate behaviors. Supporting this notion, *Grp*+ cells were predominantly suppressed by full field visual flows, whereas *Vip*+ (and *Cbln4*+) cells showed strong responses. These response patterns suggest that *Grp*+ cells are specialized for processing rapid motion caused by small objects, in line with their likely involvement in innate behaviors. *Vip*+ and *Cbln4*+ cells, each exhibiting distinct feature preferences, may be well suited for computing full field visual motion and potentially support more complex processing.

Together, our studies have discovered a genetic code of visual processing in the SC, where transcriptomic cell types encode a unique combination of visual features with their specific morphology, synaptic inputs, and projection patterns. Beyond the SC, a tight correspondence between transcriptomic and functional cell types has been shown in the retina, including the direction selective ganglion cells^43,45,63–65^. This thus raises an exciting possibility that the genetic code of retinal and SC neuron subtypes may be responsible for wiring the type-specific retinocollicular innervation as revealed by our Rabies tracing experiments, and subsequently for generating specific visual responses in the SC. In the visual cortex, however, such a close correspondence does not appear to exist for basic feature selectivity, but for more complicated properties^6,66–68^. In other words, the relationship between transcriptomic and functional properties may depend on brain areas, and consequently their developmental and evolutionary origin. We are in fact just beginning to understand how transcriptomically defined cell types relate to morphological and functional properties. Our study therefore provides valuable information to the fundamental question of “how to define a cell type” in the nervous system^2^.

## Methods

### Animals

This study used strains of Cbln4-IRES-Cre, Vip-IRES-Cre, Grp-KH288-Cre mice, Ai9, C57BL/6J mice of either sex between 6 weeks and 10 months. Five Cbln4-IRES-Cre mice were used to verify the faithfulness of Cre in labeling *Cbln4* expression in the sSC. Three Vip-IRES-Cre x Ai9 mice were used to examine the validity of this line in the sSC. Thirteen Cbln4-IRES-Cre mice, 11 Vip-IRES-Cre mice, and 11 Grp-KH288-Cre mice were used for two photon calcium imaging. Three Cbln4-IRES-Cre, 3 Vip-IRES-Cre, and 3 Grp-KH288-Cre mice were used for morphology study. Two Cbln4-IRES-Cre, 5 Vip-IRES-Cre, and 5 Grp-KH288-Cre mice were used for monosynaptic rabies virus tracing. Four Cbln4-IRES-Cre, 4 Vip-IRES-Cre, and 3 Grp-KH288-Cre mice were used to trace axon terminal expression in PBG/LGN. Seven C57BL/6J mice were used for retrograde CTB experiments. All mice were kept on a 12 h/12 h light/dark cycle in the animal room. Two to five animals were housed per cage. All experimental procedures were approved by the University of Virginia Institutional Animal Care and Use Committee.

### Generation of Cbln4-IRES-Cre knock-in mice

The Cbln4-IRES-Cre mice were generated by Augusta University Genome Editing Core (RRID:SCR_027269) using CRISPR/Cas9 approach. Briefly, the DNA repair template containing the IRES-Cre gene cassette was synthesized and purified by Integrated DNA Technologies (Coralville, Iowa 52241 USA). A synthetic guide RNA (sgRNA), which targeted the *Cbln4* C-terminal sequen5’-GGAGAAGTCGGGTTCTATAG-3′ was synthesized by Synthego (Redwood City, CA). Ribonucleoprotein (RNP) complexes were formed with sgRNA and SpCas9 nuclease 2NLS (Synthego) and injected along with the repair template into fertilized eggs from C57BL/6J mice (Jackson Laboratory, Stock #000664). Viable two-cell stage embryos were transferred to pseudo-pregnant Swiss Webster females (Taconic Biosciences, Inc., Germantown, NY, Stock #SW-F-EF) to generate founder mice. The positively targeted founder mice were identified by external long range PCR amplification of a 3,037 bp fragment containing IRES-Cre using primers 5’-GGGAGCAGCTGGTGCTATCTGACCA-3’. and 5’-GCCAAGGTTTGGCGCTGAGCA-3’ and were confirmed by Sanger sequencing by Azenta Life Sciences (Research Triangle Park, NC). Founder mice were subsequently bred with wild type C57BL/6J mice for germline transmission to generate F1 heterozygous mice that were confirmed by external long range PCR genotyping and Sanger sequencing.

### Perfusion and slicing

A lethal dose of Euthasol was intraperitoneally injected to deeply anesthetize the animals. Subsequently, transcardiac perfusion was performed with 1x PBS followed by 4% paraformaldehyde (PFA). The eyes were collected when needed, and the brains were dissected from connected tissues. The eyes were fixed in 4% PFA at room temperature for 30 mins and transferred to PBS for dissection and staining. The collected brains were kept in the dark and fixed in 4% PFA at 4°C overnight. Next, they were transferred into 30% sucrose at 4°C for 3 days for dehydration. Once sunk to the bottom of the container, brains were embedded in O.C.T. compound and frozen in a -80°C refrigerator. For slicing, brains were transferred into cryostat (Leica CM1950) at -20°C for more than half hour and coronally sliced into 40 µm sections. The brain slices were collected and stored in cryoprotectant at -20°C for subsequent imaging or staining procedures.

For morphology, brains were coronally sliced at 80 µm on the Leica VT 1000 S vibratome (Leica Biosystems). The sections were collected into four wells, stored in 0.05% sodium azide (NaN3) in 0.01 M PBS at 4 degrees Celsius, prior to mounting for confocal microscopy.

### RNA Fluorescence in situ hybridization (FISH) and immunohistochemistry

RNA FISH followed procedures described in our previous publication^32,40^. To start staining, brain sections were transferred from cryoprotectant to 1x phosphate-buffered saline (PBS, pH 7.2) at room temperature. The sections were washed twice in 1 x PBS for 10 minutes each and treated with 30% hydrogen peroxide for 10 mins to quench endogenous peroxidase activity. The bubbles on the brain sections were gently washed off by a brush in 1 x PBS. The sections were again washed in 1 x PBS three times for 10 minutes and mounted on Fisher slides (Cat. 12-550-15) to dry overnight. The following day, staining was carried out using RNAscope Multiplex v2 (Cat. 323100) reagent kit as the respective assay protocols ^69^. Briefly, slide-mounted slices were rinsed with PBS, dehydrated sequentially in 50%, 70%, and 100% ethanol, circled with a hydrophobic barrier, treated with protease IV, and then incubated with targeted probes at 40°C for 2 hours. Subsequently, they were subjected to Amp 1-3 treatments. Probes utilized included *Cbln4*, Cre, *Vip, Grp, Slc17a6 (*vGluT2), and *Slc32a1 (*vGAT). Finally, HRP-C1/C2/C3, dye FITC/Cy3/Cy5, and an HRP blocker were applied to bind Amp3 and trigger the fluorescence.

Immunohistochemistry was employed to enhance the fluorescence of helper and rabies signals as well as CTB signals. Post FISH, brain sections underwent two washes with PBST (0.5% Triton X-100 in PBS) for 5 minutes each. Sections were then incubated in blocking buffer (0.5% Triton X-100 and 5% normal donkey serum, NDS in PBS) for 1 hour at room temperature. The primary antibody (anti-GFP-chicken for helper virus and anti-RFP-rabbit for rabies, or anti-CTB-rabbit were diluted 1:300 in blocking buffer) was applied to brain sections overnight at 4°C. The following day, sections were incubated with a secondary antibody (anti-chicken 488 for helper virus and anti-rabbit 594 for rabies, or anti-rabbit 594 for CTB were diluted 1:250 in PBS) for 2 hours at room temperature. Finally, DAPI was used as a nuclear counterstain, and fluorescence images were acquired using a confocal microscope (Zeiss LSM 800) with objectives ranging from 10X/0.45 to 20X/0.80 Plan Apo.

Immunohistochemistry was also applied to stain *Cartpt* and *Spp1* in the retina to study the cell types of rabies+ retinal ganglion cells. The procedures followed the methods described above except the whole staining was performed for floating retina. The primary antibodies were anti-rabbit-Cartpt and anti-goat-Spp1. The secondary antibodies were anti-rabbit 488 for *Cartpt* and anti-goat 647 for *Spp1*.

Tissue imaged for morphology was mounted on Superfrost Plus Slide and cover slipped with Aqua Poly/Mount Mounting medium. Images were collected using Leica Stellaris 5 laser scanning confocal microscope. Images used in Fig. 5a were collected using objective HC PL APO 40×/1.3 at 4K resolution with a 246 x 246 μm field of view. Images were collected as tiled z stacks of the sSC.

### Surgery for stereotaxic injection and SC imaging

Animals were initially anesthetized with 5% isoflurane in a sealed container and transferred to a stereotaxic apparatus (Digital Model 1900, Stereotaxic Alignment System, Kopf Instruments) with a heating pad to keep the body temperature. During surgery, the animals were continuously given isoflurane (∼2% for maintenance, in O_2_, ∼0.5 L/min; VetFlo, Kent Scientific) and the concentration was adjusted based on the animal condition. The animal was loosely fixed with ear bars and a bite bar. Subsequently, the hair was cut followed by a midline incision for tracer/viral injection. The skull position was adjusted to align the Bregma and Lambda in the same plane and the head was then fixed tightly. The injection site was marked and drilled based on the coordinate of target areas. The coordinates referred to mouse brain atlas: sSC (ML: 0.8 mm; AP: −3.8 mm; DV: 1.2 mm), LGN (ML: 2.1 mm; AP: −2.3 mm; DV: 2.7 mm), PBG (ML: 1.95 mm; AP: −4.4 mm; DV: 3.1 mm). A glass pipette loaded with tracer/virus was fitted with a pressure injector (Drummond Scientific Nanoject II). A total volume of ∼100 nl virus was injected into the SC and the wait period was 3-6 weeks to verify the mouse line, trace the projection pattern, and analyze morphology. ∼50 nl CTB was injected into the LGN/PBG, and the wait period was 7 days for retrograde tracing. For rabies study, ∼200 nl helper virus was delivered into SC and followed by ∼300 nl rabies injection at the same site 3-4 weeks later. The expression pattern was examined 5-7 days after the rabies injection. After surgery, carprofen (5 mg/kg) was injected subcutaneously and the animals recovered on a heating pad before being returned to the home cage.

For two-photon imaging, animals underwent the same surgical procedure. A 2.5 mm craniotomy was made above the SC by thinning the skull with a dental drill, followed by carefully removing the skull pieces. The dura above the SC was cut to expose the posterior of SC for viral injection. 2-3 injections were made into the left hemisphere of SC, each at two depths just below the surface. ∼ 50 nl of AAV-hSyn-FLEX-GCaMP6s were delivered at each depth. After injection, a four-layer custom-made cranial window was inserted from posterior to anterior to push the sinus away from the SC and glued down using Vetbond (3M). Subsequently, a titanium head plate and ring were fixed around the imaging window using Metabond (Parkell) mixed with black ink.

### *In vivo* two photon calcium imaging

Two to three weeks after surgery, mice were habituated with the imaging setup, and GCaMP6s expression was examined. Once the labeled neurons were visually active, imaging was performed 3-6 weeks after surgery in awake head-fixed mice placed on a cylindrical Styrofoam treadmill. Neuronal activity was recorded by a two-photon scanning microscope (Ultima Investigator, Bruker Nano Surface Division) equipped with a Ti:sapphire laser (Chameleon Discovery with TPC, Coherent) and a 16×, 0.8 NA Nikon objective at the laser wavelength of 920 nm. PrairieView software (version 5.8) was used to control laser power, optical zoom, imaging depth, and resolution. It also simultaneously recorded the timing of the visual stimulus through a third channel (other than red and green channels). In this study, imaging data were acquired using resonant scanning mode at 2.5x optical zoom, 30 Hz, and 512 x 512 pixel resolution. The images were averaged every four frames for later analysis. To image the soma of the 3 cell types, *Cbln4*+ and *Vip*+ cells were imaged at ∼60 µm and *Grp*+ cells were imaged at ∼230 µm below SC surface. An animal was recorded only once (one session) and no more than 2 hours when the required laser power was low (∼35 mW), and no more than 1 hour when laser power was high (>70 mW). Animals were free from experiment for at least two days after each imaging session.

### Visual stimulus

An LCD monitor (52.7 x 29.6 cm, Dell, SE2422HX, 60 Hz refresh rate, gamma corrected) was used to display the visual stimulus to the animal’s right eye (contralateral to the GCaMP6s injection and imaging site). MATLAB Psychophysics toolbox^70^ was used to generate all the stimuli as described in our previous studies, except ‘flows’ were generated using a custom Python script and presented by PsychoPy^71^. Before each imaging session, a 10° black square was flashed on a gray background, 0.5s On and 0.5s Off, from up to down and left to right sequentially on the monitor. Only sessions in which GCaMP6s+ neurons showed clear activity progression following the position of flashing squares were recorded and included for further data analysis. The monitor was accordingly adjusted to match the receptive field of the GCaMP6s+ cells at the center of the screen, which was 25 cm away from the right eye. The receptive field of almost all imaged neurons was covered by the stimuli used in our studies. All stimuli were centered on the screen with gray background and repeated 10 times in a pseudo-randomized sequence unless otherwise stated.

10° black flashing squares were used to map and measure the receptive field. The squares were displayed at a 5 by 5 grid (10° spacing). Each flash was displayed for 1s with a 1s interval.

Sinusoidal drifting gratings with certain speeds were used to study direction selectivity. A fast grating was displayed at SF of 0.08 cpd and TF of 2 Hz (25°/sec, 1s on and 4s off). Two slow gratings were displayed either at SF of 0.16 cpd and TF of 0.8 Hz (5°/sec, 1s on and 4s off) or SF of 0.08 cpd and TF of 0.5 Hz (6.25°/sec, 2s on and 4s off). All the gratings were displayed at a 40° (diameter) patch covering the receptive fields, in 100% contrast and 12 different moving directions (0° to 330°, 30° spacing). 0° direction was for nasally directed motion and 90° for upward motion on the stimulus monitor.

In addition, sinusoidal drifting gratings with a series of SF (0.02-0.64 cpd, 6 logarithmic steps) and TF (0.25-8 Hz, 6 steps) combinations were presented to study the spatial and temporal preference. The stimulus was displayed at 90° for *Cbln4*+ and *Grp*+ cells and 330° for *Vip*+ cells. The gratings were also presented at a 40° patch in diameter. Each condition was displayed for 4s with 4s inter-stimulus interval.

Black flashing squares of different sizes (2° to 64°, 6 steps) were used to measure cell’s size preference. The squares were presented in the center of the screen with 1s on and 4s off. This was done in the same session as receptive field mapping so that we could determine the receptive field of the imaged neurons. Only those with the receptive field in the center of the screen were selected to analyze size tuning.

Black “looming” disks of different expanding speeds (5 °/s to 160 °/s, 6 steps; expanding from 0 to 60° diameter) were used to study speed tuning. Inter-stimulus interval was 4s. For this stimulus set, 10 repeats were recorded, except in one Vip-IRES-Cre mouse (n = 12 *Vip*+ cells; 6 repeats).

“Flows” were presented following the design by Dyballa et al. (2018)^57^, which used continuous, non-rigid and locally correlated flow patterns to provide a more naturalistic representation of visual motion while retaining precise experimental parameter control. Specifically, motion generation incorporated Boids-inspired collective-motion rules^72^ to produce smooth flow-like trajectories. The stimuli consisted of a full-field pattern of black dots and were parametrically varied in dot size (diameter, 2°, 4°, 8°, 16°), speed (0, 5, 10, 20, 40 °/s), and motion direction (12 directions). Short-range separation forces maintained a relatively uniform spatial distribution of dots and prevented local clustering. A local direction field with fixed directional dispersion (motion jitter: standard deviation = 0.09) to produce smooth, continuous, and locally correlated flow-like motion. Each condition was presented for 1s, followed by a 2s gray inter-stimulus interval. All conditions were presented randomly and repeated five times.

### Analysis of RNA FISH and immunohistochemistry

The quantification of probe-positive and Cre+/rabies+/CTB+ cells was performed using Zen 3.4 software (blue edition), based on puncta density for FISH and fluorescence intensity for tdTomato. First, the regions of interest were selected as follows: (1) For mouse line verification, both Cbln4-IRES-Cre mice and Vip-IRES-Cre mice, a 250 µm * 150 µm rectangle was drawn from the SC surface for each SC image. (2) For rabies+ cells in the retina, each image was acquired at a 319.45 µm * 319.45 µm square. (3) For rabies+ cells in the sSC, two 51789.37 or 51645.36 µm^2^ squares were randomly drawn near the surface of each SC image. (4) For CTB+ cells in the sSC, a square of 19533.49 µm^2^ was drawn in the area with good expression of both CTB and marker genes.

Second, positive neurons were manually labeled by the Points Events function in the Zen software for each selected region. Probe-positive cells were selected when a group of puncta showed a neuronal shape, and the puncta density was higher than the background noise. Similarly, Cre+/rabies+/CTB+ cells were marked when the fluorescence intensity of a cell was higher than the background. When both conditions were met and they shared similar size and shape, the cells were considered as colocalized cells.

Specificity and sensitivity were calculated as: specificity = Number of cells positive to both transcript and Cre / Number of Cre+ cells; sensitivity = Number of cells positive to both transcript and Cre / Number of transcript+ cells. Rabies+ or CTB+ percentage was calculated as Number of cells positive to both probes and rabies (CTB) / Number of rabies+ (CTB+) cells.

### Analysis of two-photon calcium imaging data

Processing of imaging data followed our published procedures to calculate ΔF/F ^32,73^. Briefly, a pre-processing procedure written in MATLAB was used to correct image shift caused by motion artifact. Regions of interest (ROIs) of individual neurons were manually drawn on the average image over the entire time-series or with clear responses from viewing the recording. To compare the response of same cells to different stimuli, ROIs were selected for cells identified in multiple recording sessions. The fluorescence magnitude ΔF/F_0_ was calculated from (F − F_0_)/F_0_, where F_0_ was the mean of all pixels in each ROI over six frames before the stimulus onset, and F was the average fluorescence signal over the duration of stimulus presentation, starting from one frame after the stimulus onset. The ΔF/F_0_ was further averaged across all repeats for each cell and used for further analysis.

To identify whether a cell was responsive, three criteria needed to be met. First, the maximum ΔF/F_0_ was > 0.2 to at least one stimulus condition. Second, the mean ΔF/F_0_ was larger than 2 SDs above the baseline signal for at least one condition. Third, the bootstrap test examining trial-by-trial variability was significant for at least one condition. Specifically, the stimulus evoked ΔF/F_0_ and the response of blank stimulus condition (or the pre-stimulus baseline) of each repeat of a condition were pooled with replacement over 10000-iteration resampling. The difference between the mean of the resampled stimulus and control response was calculated for each iteration. A response condition falling outside the 95% confidence interval of the resampled null distribution considered significant.

The “center-of-mass” was calculated to determine the receptive field center^37^: 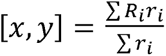 where *i* is the location in the grid at which the cell was responsive. *R* and *r* are ΔF/F_0_ and position vector at the th location, respectively. To measure receptive field size, the number of responsive subfields in the grid was counted and multiplied by the square size (10° x 10°). The receptive field size along x and y axis was further calculated by the responses (ΔF/F_0_) with a two-dimensional Gaussian distribution^37^. Only neurons with good fitting (r^2^ > 0.7) were included to plot the distribution of x and y sizes.

The preferred direction was the angle of vector sum of ΔF/F_0_ cross 12 directions. The global direction selective index (gDSI) was calculated as the vector sum of ΔF/F_0_ normalized by their scalar sum. The global orientation selective index (gOSI or called global axis selective index: gASI) was calculated similarly except doubling the angle in the equation (thus folding the responses of opposite directions onto the same orientation to 0 - 180°). Both calculations generate a value between 0 and 1, and 0.2 was used as the cutoff to determine direction selective cells or orientation/axis selective cells. The equations are shown below:

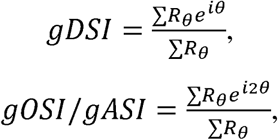

where is ΔF/F_0_ at each direction of the stimulus.

The tuning width for drifting gratings was calculated as the full width at half-height (FWHM) for responsive cells. The responses at 12 stimulus directions were fit with a sum of two Gaussians to obtain a smooth tuning curve at the resolution of 1° over 360°. The FWHM was further analyzed for cells with R^2^ > 0.7 from the smooth tuning curve. FWHM was measured only for the main peak if a cell showed two-peak tuning such as most *Grp*+ cells.

To plot tuning curves for flow, the response magnitude to static dots of the same size was subtracted for each neuron, leaving the response only evoked by the moving dots of certain size, speed, and direction. Therefore, the response magnitude formed a 3D matrix with 192 conditions including 12 directions x 4 sizes x 4 speeds. The “overall” tuning curve for a particular feature (e.g., direction) was the average of all responses over the other two dimensions (e.g., average of 4 sizes and 4 speeds at each direction). To determine the tuning curve at the “best” condition, the max response was identified from all 192 conditions, and the corresponding combination of features was considered the preferred direction, size and speed. The tuning curve for a specific feature (e.g., direction) was plotted using responses at the preferred condition of the other 2 features (e.g., at the preferred size and speed).

### Analysis of encoding manifold

To characterize population-level functional organization, flow-evoked responses of 380 neurons (184 *Cbln4^+^*, 81 *Vip^+^*, and 115 *Grp+* neurons from 4, 5, and 5 mice, respectively) were embedded into a shared encoding manifold^41^. For each neuron, Δ*F/F* signals were averaged over stimulus time and across five repetitions per stimulus to construct a 196-dimensional mean response vector, which was subsequently mean-subtracted across conditions and normalized to unit norm to isolate relative tuning profiles. Dimensionality reduction via PCA retained 102 components accounting for at least 95% of the variance.

Within this 102-dimensional PCA space, a symmetric -nearest-neighbour graph (*K*=15, Euclidean distance) was constructed, connecting neuron pairs whenever either neuron ranked among the other’s 15 nearest neighbors. Pairwise affinities were weighted using a locally scaled Gaussian kernel, with each neuron’s local scale set to its 15th-neighbour distance^74^. Following anisotropic density normalization (α=1) to correct for non-uniform sampling density, the affinity matrix was row-normalized into a Markov transition matrix. Spectral decomposition via diffusion maps^41,75^ extracted the top three non-trivial eigenvectors as a 3D manifold embedding. Cell-type identities and preferred stimulus parameters (determined from the condition evoking the peak unnormalized mean response) were mapped *post hoc* onto the fixed embedding coordinates.

Inter-animal consistency and stimulus-feature contributions were evaluated in the full normalized 196-dimensional space. Pairwise cosine similarities and Euclidean distances were averaged across neuron pairs within each animal pair before taking the unweighted mean across animal pairs. Neighborhood coverage between animal pairs was quantified as the fraction of source-animal neurons having at least one target-animal neuron among their 15 nearest neighbors.

### Analysis of neuron morphology

Neuron morphology was assessed using a total of seven morphometric variables extracted from volumetric confocal datasets of the imaged mouse lines based on previous work^29^. These variables were: soma volume, soma distance to SC surface, total number of Sholl intersections, number of primary dendrites, dendritic arbor orientation index, maximum number of intersections, radii at which the maximum number of intersections occur.

For dendritic measurements, z-stacks were imported to Fiji (ImageJ) and the neurites were reconstructed using the SNT Plugin (5.0.11) as detailed previously^49^. Briefly, each cell was visualized by semi-manually tracing individual dendrites as “paths”. Starting from the soma each primary dendrite was traced until its branching point in which the secondary paths were traced as “daughter paths” of the primary dendrite. Traces were then filled, creating a binary mask and finalized with the “Close-” function to limit holes in the mask. Each mask was maximum projected to eliminate the z-depth. The binary images were rotated manually to account for the angle of the dorsal SC surface before proceeding with further data collection. Two straight lines were overlaid on the cell, one parallel and one perpendicular to the SC surface to measure the horizontal and the vertical dendritic arbor width respectively (Extended Data Fig. 5a). In SNT, the rotated binary masks were used to run a Sholl analysis to investigate dendritic orientation and expansiveness. The Sholl analysis plugin was used at 15 µm intervals to capture Total Sholl Points, Maximum Intersections and Radii of Maximum Intersections.

The dendritic arbor orientation index (DAOI) normalized each neuron and calculates the orientation of dendrites. Each image was parceled into 8 quadrants originating from the center of the soma. Each quadrant was 45 degrees and is classified as either vertical (315-45, 135-225) or horizontal (225-315, 45-135) quadrants using a custom macro script (See Data and code availability: https://doi.org/10.18130/V3/WTVJYB). The parceled images were individually analyzed with the SNT Sholl Analysis to count the Total Sholl Points. The following equation was used to calculate Dendritic Arbor Orientation Index. A value between –1 to +1 was calculated with +1 being a completely vertical neuron and –1 a completely horizontal neuron.

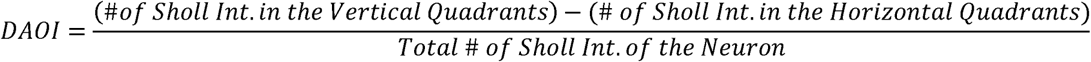

For somatic measurements, z-stacks were initially imported to Imaris 10.2.0 (Oxford Instruments) where soma were segmented with a pixel classifier, converted into 3D objects and assigned unique identifiers. SC surface was manually delineated as a separate object and then both the soma volumes (µm3) and the shortest distances (of each soma to the SC surface) (µm2) were extracted using the object-object statistics within the statistics panel.

All values reported were mean ± SEM unless otherwise mentioned. The number of replicates and statistical tests are listed in the figure legends.

## Acknowledgement

We thank Dr. Lin Gan for generating the Cbln4-IRES-Cre mice; Dr. David Olson for providing the Vip-IRES-Cre mice; Dr. Martha Bickford for providing the Grp-KH288-Cre mice. This work was supported by US NIH grants (EY026286 and EY020950) and Jefferson Scholars Foundation endowment to J.C.

## Author Contributions

J.C. and C.C. designed the experiments; C.C. verified the mouse lines; C.C. performed two-photon imaging experiments, with assistance of surgery from C.L. and data analysis from H.C. J.R. programmed the flow stimuli and performed encoding manifold analysis. D.E. performed morphology experiments, with assistance of surgery from C.C. and data analysis from A.K. and A.E. C.C. and Y.L. did surgery and collected brain sections or tracing experiments; Y.L. dissected the retina and performed RNA FISH and immunohistochemistry; C.C. performed confocal imaging and the quantifications. C.C. and J.C. wrote the manuscript with input and approval from all authors.

## Data availability statement

The data supporting the findings of this study are available from the corresponding author upon reasonable request.

