## Supplemental Figures for "Distinct processing of visual motion by transcriptomic cell types in the superior colliculus"

a

| Type | Stimulus | Conditions | Animal | Total cells | Responsive cells | Proportion |
|---|---|---|---|---|---|---|
| *Cbln4* | RF map | 25 | 5 | 159 | 102 | 64.15 |
| | Size | 6 | 4 | 26 | 21 | 80.77 |
| | DG-25d/s | 12 | 5 | 323 | 192 | 59.44 |
| | DG-5d/s | 12 | 6 | 269 | 182 | 67.66 |
| | SFTF | 36 | 4 | 186 | 172 | 92.47 |
| | Looming | 6 | 7 | 253 | 194 | 76.68 |
| | Flow | 196 | 4 | 184 | 168 | 91.30 |
| *Vip* | RF map | 25 | 8 | 117 | 80 | 68.38 |
| | Size | 6 | 8 | 20 | 18 | 90.00 |
| | DG-25d/s | 12 | 6 | 88 | 21 | 23.86 |
| | DG-5d/s | 12 | 7 | 107 | 75 | 70.09 |
| | DG-6d/s | 12 | 4 | 38 | 37 | 97.37 |
| | SFTF | 36 | 3 | 23 | 23 | 100.00 |
| | Looming | 6 | 6 | 92 | 73 | 79.35 |
| | Flow | 196 | 5 | 81 | 80 | 98.77 |
| *Grp* | RF map | 25 | 7 | 139 | 123 | 88.49 |
| | Size | 6 | 5 | 32 | 32 | 100.00 |
| | DG-25d/s | 12 | 9 | 232 | 119 | 51.29 |
| | SFTF | 36 | 3 | 56 | 45 | 80.36 |
| | Looming | 6 | 8 | 211 | 141 | 66.82 |
| | Flow | 196 | 5 | 115 | 33 | 28.70 |

b

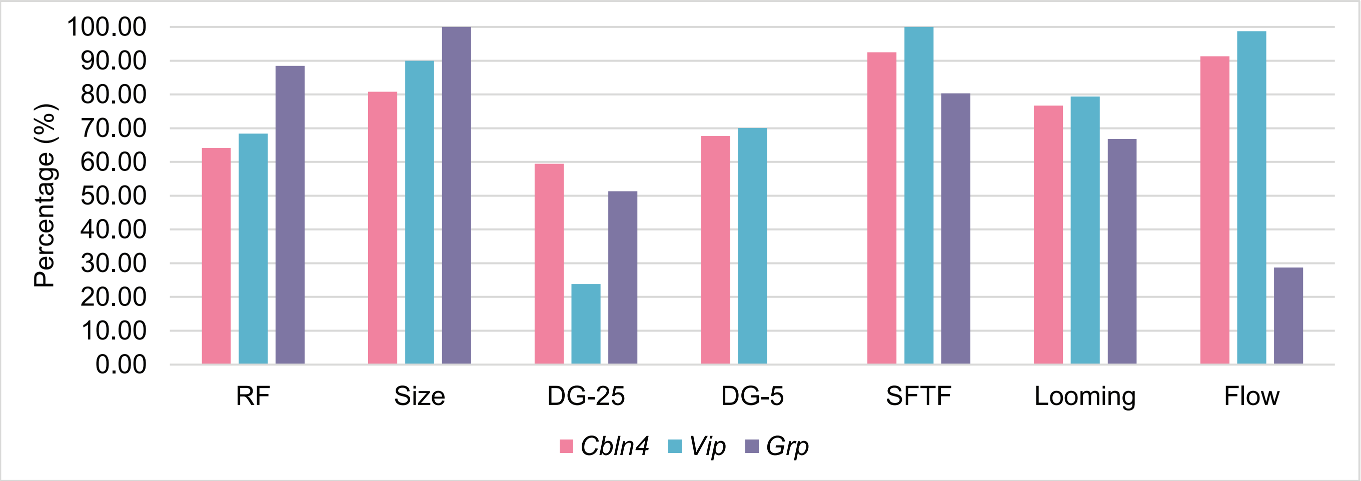


**Extended Data Fig. 1 Summary of animal numbers, cell numbers, and responsiveness of 2-p imaging experiments**

**a,** Summary table of 2-p recordings of three cell types to different stimuli.

**b,** Bar plot showing the proportion of responsive cells of three cell types to different stimuli.

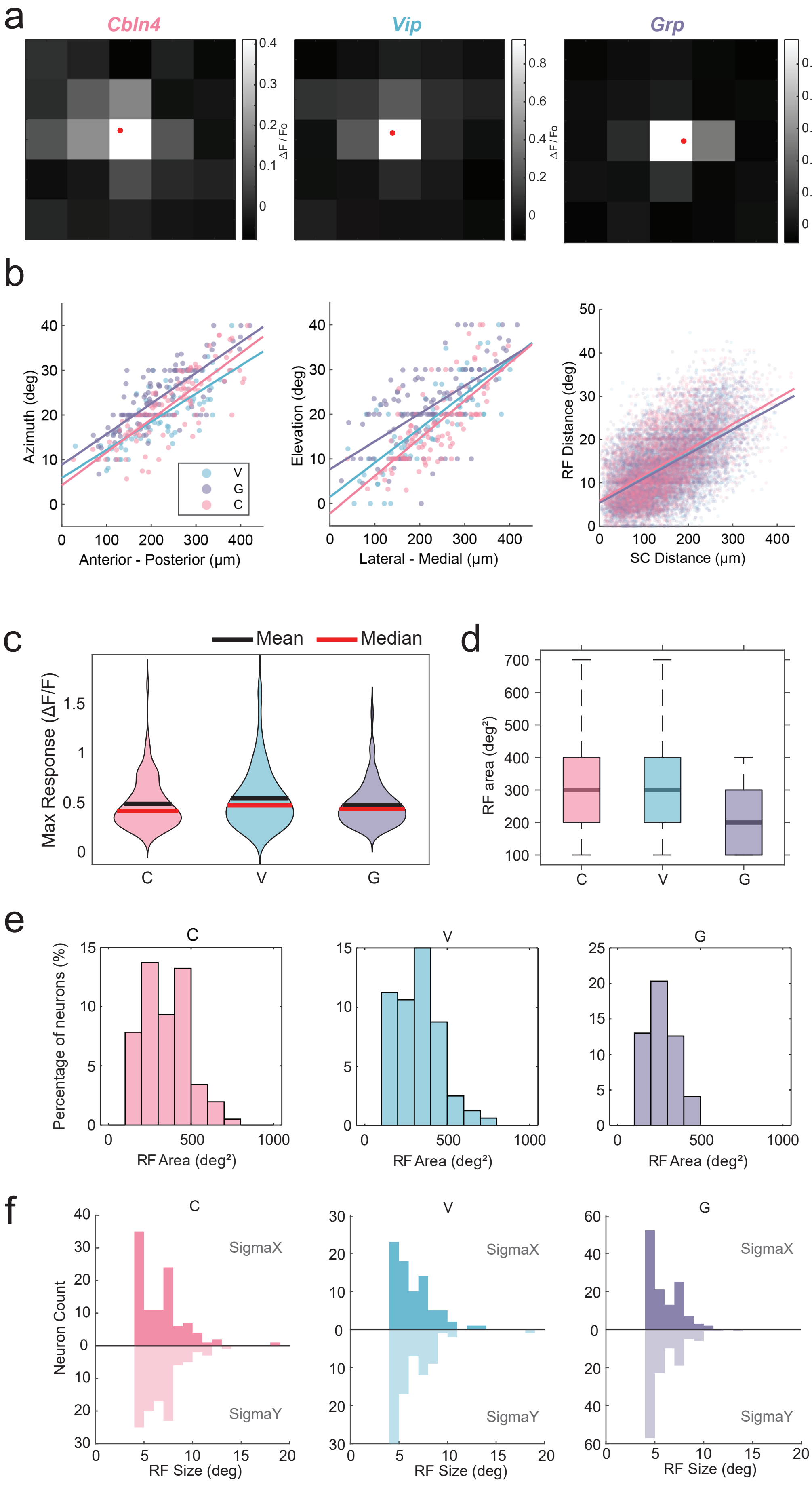

a
Cbln4
Vip
Grp
ΔF / Fo
b
Azimuth (deg)
Anterior - Posterior (µm)
V
G
C
Elevation (deg)
Lateral - Medial (µm)
RF Distance (deg)
SC Distance (µm)
c
Mean
Median
Max Response (ΔF/F)
C
V
G
d
RF area (deg²)
C
V
G
e
C
V
G
Percentage of neurons (%)
RF Area (deg²)
f
C
V
G
Neuron Count
SigmaX
SigmaY
RF Size (deg)

**Extended Data Fig. 2 Receptive field mapping**

**a,** Heatmap presenting the response of example cells responding to flashing squares in different locations. Red dot: receptive field center.

**b,** Scatter plots showing the correlation between cell pair distance in the SC and receptive field distance on the stimulus monitor.

**c,** Violin plot showing the distribution of max response of the 3 cell types responding to flashing squares. C: n = 102 cells, 5 mice; V: n = 80 cells, 8 mice; G: n = 123 cells, 7 mice.

**d-e,** Box plot and histogram visualizing the distribution of receptive field size of the 3 cell types.

**f,** Histogram showing the distribution of receptive field size of x and y separately of the 3 cell types.

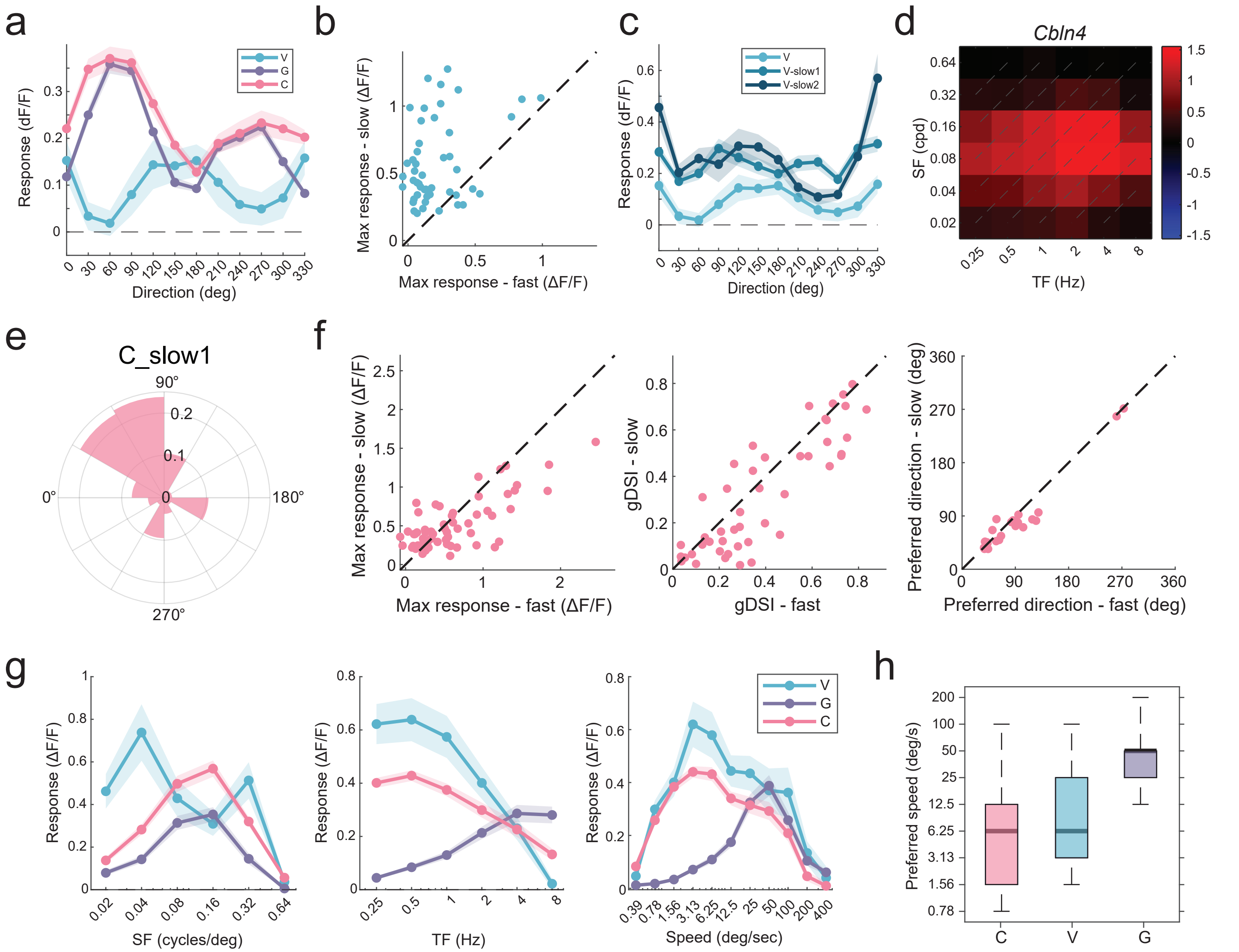


**Extended Data Fig. 3 Distinct feature preference of three cell types**

**a,** Average direction tuning curves responding to the "classic" drifting gratings (SF = 0.08 cpd; TF = 2 Hz; 12 directions. C: n = 192 cells, 5 mice; V: n = 21 cells, 6 mice; G: n = 119 cells, 9 mice. Mean ± SEM.

**b,** Scatter plot showing the max response of same *Vip*+ cells responding to the classic drifting gratings and slow drifting gratings (SF = 0.16 cpd, TF = 0.8 Hz). n = 77 cells, 5 mice.

**c,** Average tuning curves of *Vip*+ cells responding to three drifting gratings (V: SF = 0.08 cpd, TF = 2 Hz; V_slow1: SF = 0.16 cpd, TF = 0.8 Hz; V_slow2: SF = 0.08 cpd, TF = 0.5 Hz). V: n = 21 cells, 6 mice; V_slow1: n = 75 cells, 7 mice; V_slow2: n = 37 cells, 4 mice. Mean ± SEM.

**d,** Heatmap showing the response magnitude of an example *Cbln4*+ cells for different SF-TF combinations.

**e,** Rose histogram presenting the distribution of preferred direction of *Cbln4*+ cells responding to slow1 drifting gratings. n = 269 cells, 6 mice.

**f,** Comparison of the same *Cbln4*+ cells responding to drifting gratings of 2 different speeds ("fast" - SF = 0.08 cpd, TF = 2 Hz; and slow - SF = 0.16 cpd, TF = 0.8 Hz). Max response for responsive cells to either stimulus, n = 62 cells, 2 mice. gDSI for responsive cells to both stimuli, n = 46 cells, 2 mice. Preferred direction for responsive and DS cells (gDSI > 0.2) for both stimuli, n = 23 cells, 2 mice.

**g,** Average tuning curves of 3 cell types responding to SF-TF combinations. C: n = 172 cells, 4 mice; V: n = 23 cells, 3 mice; G: n = 45 cells, 3 mice.

**h,** Box plot showing the distribution of preferred speed responding to SF-TF combinations.

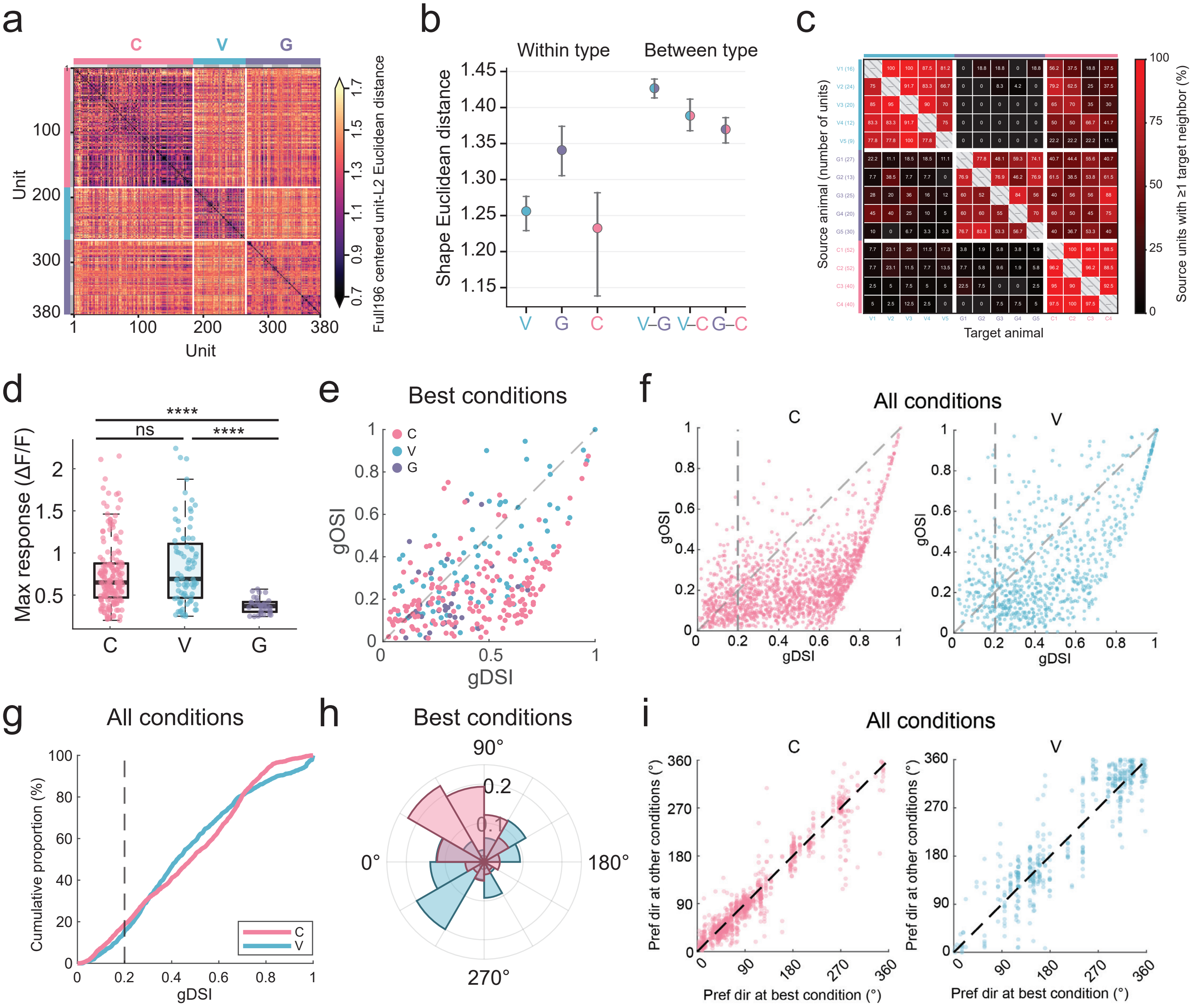


**Extended Data Fig. 4 Distinct responses to visual flows**

**a,** Heatmap presenting the pairwise Euclidean distance between all recorded neurons from the 3 cell types.

**b**, Mean shape Euclidean distance between neurons within each cell type (left) and across types (right). Mean ± SEM.

**c**, Heatmap showing the percentage of units from each source animal (rows) with at least 1 neighbor from other animals (columns) in the encoding manifold, organized by cell types.

**d**, Box plot showing the max response of responsive cells for the 3 cell types. C: n = 168 cells, 4 mice; V: n = 80 cells, 5 mice; G: n = 33 cells, 5 mice. Group compared by Kruskal-Wallis test ($H(2) = 44.03$, $p < 0.001$), with Dunn's post-hoc test (Holm-corrected). C vs V: $p = 0.239$, C vs G & V vs G: $p < 0.0001$.

**e**, Scatter plot presenting gDSI vs gOSI of individual cells at their preferred conditions. C: n = 168 cells, 4 mice; V: n = 80 cells, 5 mice; G: n = 33 cells, 5 mice.

**f**, Scatter plots showing gDSI vs gOSI of all responsive conditions. C: n = 1634 conditions, 168 cells, 4 mice; V: n = 815 conditions, 80 cells, 5 mice.

**g,** Cumulative percentage of gDSI for all responsive conditions of *Cbln4*+ and *Vip*+ cells.

**h,** Rose histogram showing the distribution of preferred directions of *Cbln4*+ and *Vip*+ cells at their best conditions.

**i,** Scatter plot comparing the preferred direction between the best condition and other DS conditions of the same cells. C: n = 1084 conditions, 168 cells, 4 mice; V: n = 548 conditions, 80 cells, 5 mice.

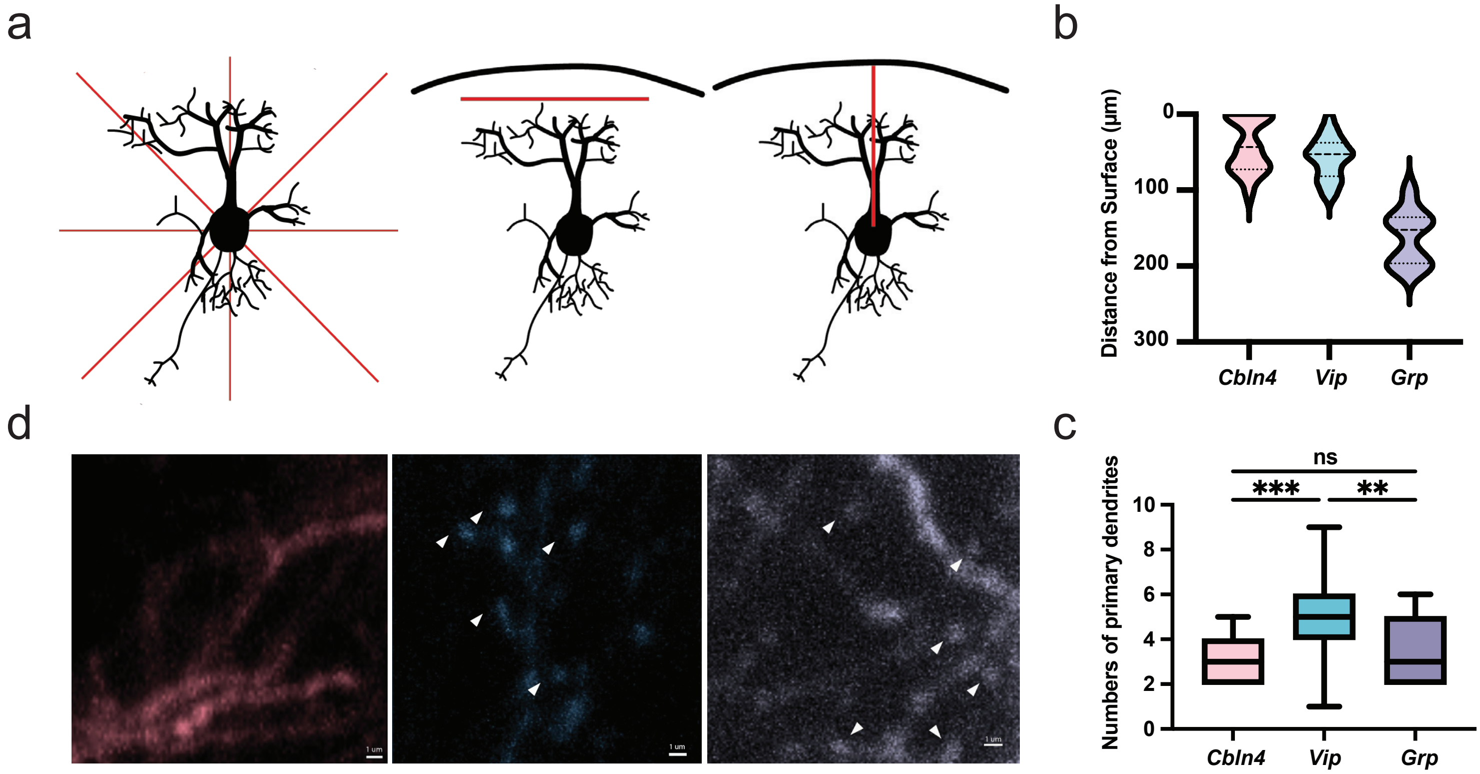


e

| Mouse Line | Soma Volume | Dist to Surf | Sholl Points | Prim Dend | DAOI | Max Int | Radii Max Int |
|---|---|---|---|---|---|---|---|
| ***Cbln4*** | 0.3015 | <0.05 | <0.05 | <0.05 | <0.05 | 0.3101 | <0.05 |
| ***Vip*** | 0.7008 | 0.8475 | 0.8106 | 0.324 | <0.05 | 0.1148 | <0.05 |
| ***Grp*** | 0.3055 | 0.1626 | 0.3861 | <0.05 | 0.0551 | 0.1692 | <0.05 |

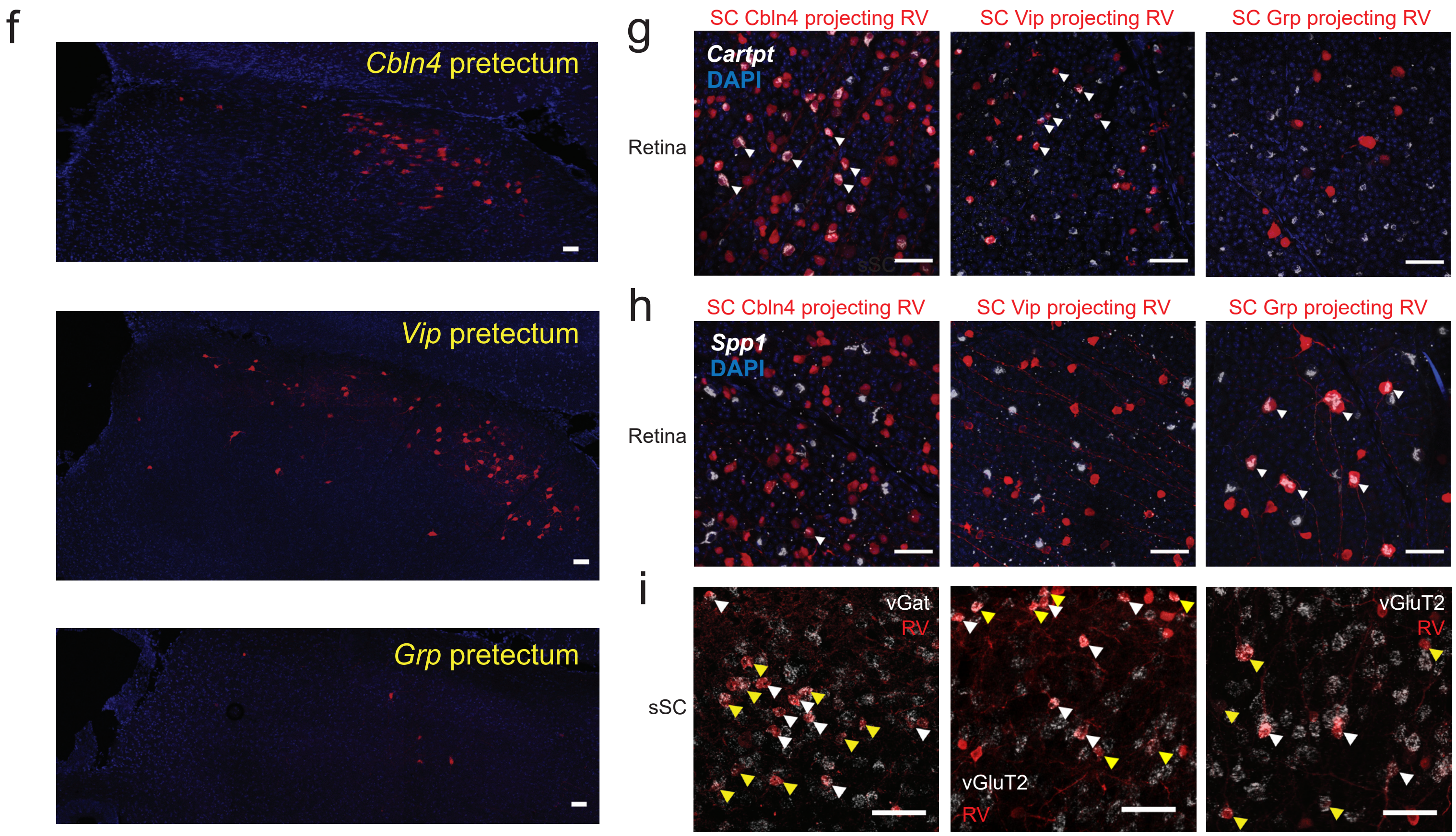

**Extended Data Fig. 5 Distinct morphology and input cell types**

**a,** Schematic illustrating quantified parameters of morphology.

**b,** Violin plot of the depth information of reconstructed cells. C: n = 20 cells, 3 mice; V: n = 23 cells, 3 mice; G: n = 20 cells, 3 mice.

**c,** Box plot showing the number of primary dendrites. C: n = 20 cells, 3 mice; V: n = 23 cells, 3 mice; G: n = 20 cells, 3 mice. Group compared by Kruskal-Wallis test ($H(2) = 18.67$, $p < 0.001$), with Dunn's post-hoc test (Holm-corrected). C vs V: $p = 0.0002$, C vs G: $p = 0.9999$, V vs G: $p = 0.0024$.

**d,** Confocal images showing dendritic spines. Example spines are labeled by white arrows. Scale bar: 1 µm.

**e,** Summary table showing Shapiro Wilks normality test for the 3 cell types based on 7 criteria: soma volume, distance to surface, sholl points, primary dendrites, dendritic arbor orientation index, maximum intersections, radii of maximum intersections. $p > 0.05$ indicates a normally distributed data set.

**f,** Confocal images presenting rabies+ expressing cells in the pretectum for the three cell types. Scale bars: 50 µm.

**g,** Example confocal image showing immunohistochemistry staining of *Cartpt* in the retina and rabies+ expressing RGCs in the 3 mouse lines. white arrow: co-localized example cells. Scale bar: 50 µm.

**h,** Example confocal image showing immunohistochemistry staining of *Spp1* in the retina and rabies+ expressing RGCs in the 3 mouse lines. white arrow: co-localized example cells. Scale bar: 50 µm.

**i,** Example confocal image showing immunohistochemistry staining of vGAT or vGluT2 in the sSC and sSC non-starter rabies+ cells. white arrow: starter cells. yellow arrow: vGAT or vGluT2+ and rabies+ non-starter cells. Scale bar: 50 µm.
